# Dynamic mRNA stability changes buffer transcriptional activation during neuronal differentiation and are regulated by RNA binding proteins

**DOI:** 10.1101/2023.09.22.558981

**Authors:** Yuan Zhou, Sherif Rashad, Teiji Tominaga, Kuniyasu Niizuma

## Abstract

The steady state levels of mRNA are outcomes of a finely tuned interplay between RNA transcription and decay. Therefore, the modulation of RNA stability is generally assumed to influence RNA abundance in a positive direction. However, the correlation between mRNA transcription, translation and stability remains elusive. Here, we employed a newly developed simplified mRNA stability profiling technique to explore the role of mRNA stability in SH-SY5Y neuronal differentiation model. Transcriptome-wide mRNA stability analysis revealed neural-specific RNA stability kinetics, including stabilization of transcripts encoding regulators of neuronal morphogenesis and function and destabilization of mitochondrial electron transport and redox homeostasis. When we further examined the relationship between transcription, translation and mRNA stability, a bidirectional regulation of RNA stability was revealed, wherein mRNA stability could either exert the buffering effect on gene products or change in a same direction as transcription. Motif analysis unveiled SAMD4A as a major regulator of the dynamic changes in mRNA stability observed during differentiation. Knockdown of SAMD4A impaired neuronal differentiation and influenced the response to oxidative stress. Mechanistically, SAMD4A was found to alter the stability of several mRNAs to which it binds. Meanwhile, a dimorphic pattern of the correlation between gene expression and SAMD4A-regulated mRNA stability was observed, suggesting dynamic regulation mRNA stability during the neuronal differentiation guided by SAMD4A. The novel insights into the interplay between mRNA stability and cellular behaviors provide a foundation for understanding neurodevelopmental processes and neurodegenerative disorders and highlights dynamic mRNA stability as an important layer of gene expression regulation.

## Introduction

Neuronal differentiation is a complex multistep process in which neurons undergo dramatic morphological alterations, including neurite outgrowth and synapse formation^1^. A series of changes then enables neuron to carry out their desired activities such as, electrophysiological activity and neurotransmitter secretion^2,3^. Consequently, these changes render mature neurons highly sensitive to oxidative stress, a property which plays a crucial etiology in many neurological and neurodegenerative diseases^4,5^. An extensive body of studies have characterized transcriptional adaptations associated with neuronal differentiation processes ranging from cell-fate commitment^6^ to synapse formation^7^. Nevertheless, the reliance only on transcriptome profiling may not fully capture the complexity of regulatory events and explain neuron-specific stress response. Recent advances highlighted the significance of mRNA decay in nervous system development, yet an understanding of the regulation of mRNA stability transcriptome-wide during neuronal differentiation and especially during neuronal stress remains elusive^8^.

RNA stability (also referred as RNA decay, RNA turnover, or RNA half-life), characterized by its variability and tight regulation on gene products, was gradually recognized as a crucial element of post-transcriptional regulation^9^. Despite substantial knowledge of the major pathways and enzymatic complexes responsible for mRNA degradation in both bacteria and eukaryotes, we still do not understand how RNA decay interplay with transcription and translation. Many studies have previously shown that RNA stability correlates with RNA levels in an organism-specific manner. More specifically, a significant positive correlation between RNA stability and RNA abundance was observed in *S. pombe*, and *S. cerevisiae* but not in *E. coli*^10^. On the other hand, recent studies have established causal links between RNA stability and translation, suggesting that these processes are intricately coupled^11^. However, to what extent this interaction pattern between RNA stability and abundance obtained in yeasts and prokaryotes can be extrapolated to human cells remains largely unclear. In addition, whether transcriptome-wide mRNA stability can dynamically impact mRNA levels and translation in various conditions is unknown.

Multiple mechanisms determine mRNA half-life^12^. One common mechanism involves the recognition of cis-elements by RNA-binding proteins (RBPs). RBP-mRNA interactions can activate or inhibit mRNA decay by affecting the recruitment or activity of RNA degradation complexes. Additional mRNA decay mechanisms include targeting by microRNAs and the nonsense-mediated decay (NMD) pathway. These mechanisms establish mRNA decay networks in which mRNA stability is genetically programmed, tunable, and tightly regulated. It was reported that mRNA decay network regulated by RBP Pumilio contributes to the relative abundance of transcripts involved in cell-fate decisions and axonogenesis during Drosophila neural development^13^. Moreover, the family of SRSF (Serine-rich splicing factor) and ELAV (Embryonic lethal abnormal vision) were successively described as the essential factor in manipulating the development and maturation of neurons^14,15^. In addition to the phenotype of differentiation, emerging evidence unveiled a stress-associated mRNA stabilization in bacteria and a strong link between altered mRNA stability and neurodegenerative disease^16^. This evidence highlights the need to discover novel RBPs of neuronal differentiation and further mechanistically understand their role.

In this study, we present a simplified method to analyze transcriptome-wide differential mRNA stability changes in dynamic conditions. We applied this method to SHSY-5Y neural differentiation model. Our data show that mRNA stability dynamically fine-tunes mRNA levels and translation by acting as a buffer between these processes. Our results also reveal SAMD4A (Sterile alpha motif domain containing protein A) as a major regulator of mRNA stability and neuronal differentiation as well as neuronal responses to oxidative stresses.

## Result

### ➢ Neuronal phenotypic characterization of differentiated SH-SY5Y cells

We observed the neuronal characteristics of differentiated SH-SY5Y cells from the two aspects: neuronal morphological and biochemical changes and neuronal response to oxidative stress. SH-SY5Y cells were subjected to 10-days differentiation treatment according to the protocol in **Figure 1A**. Undifferentiated SH-SY5Y cells grew in clumps and demonstrated a large, flat, epithelial-like cell body with numerous short processes extending outward^17^, while pre-differentiated cells by 4-day RA treatment exhibited disassociated clump, cell body shrinkage and possessed several neurites outgrowth, and fully-differentiated cells by subsequent 6-day BDNF treatment developed a triangle cell body and extensive neurites that projected to surrounding cells to form complex neurite network (**Figure 1B**). **Figure 1C** demonstrated the differential expression of all the neuronal markers (MAP2, TUJ and SYN) in the Diff. compared to the Undiff. cells. It was noted that the expression of MAP2, TUJ and SYN in Diff. group followed a polarized distribution into growing neurites, a distinctive feature of mature neurons^18^. Moreover, western blot analysis on the protein expression level of neuronal markers and stem cell markers showed that the significantly lower expression of the immature neuronal marker Nestin and stem cell markers SOX2 and Nanog, and higher expression of mature neuronal markers MAP2, NF-M, PSD95, TUJ and SYN in the Diff. cells than Undiff. cells (P < 0.05; **Figure 1D&E**). Taken together, these data validate the success in differentiating SHSY-5Y into neuron-like cells.

**Figure 1.**
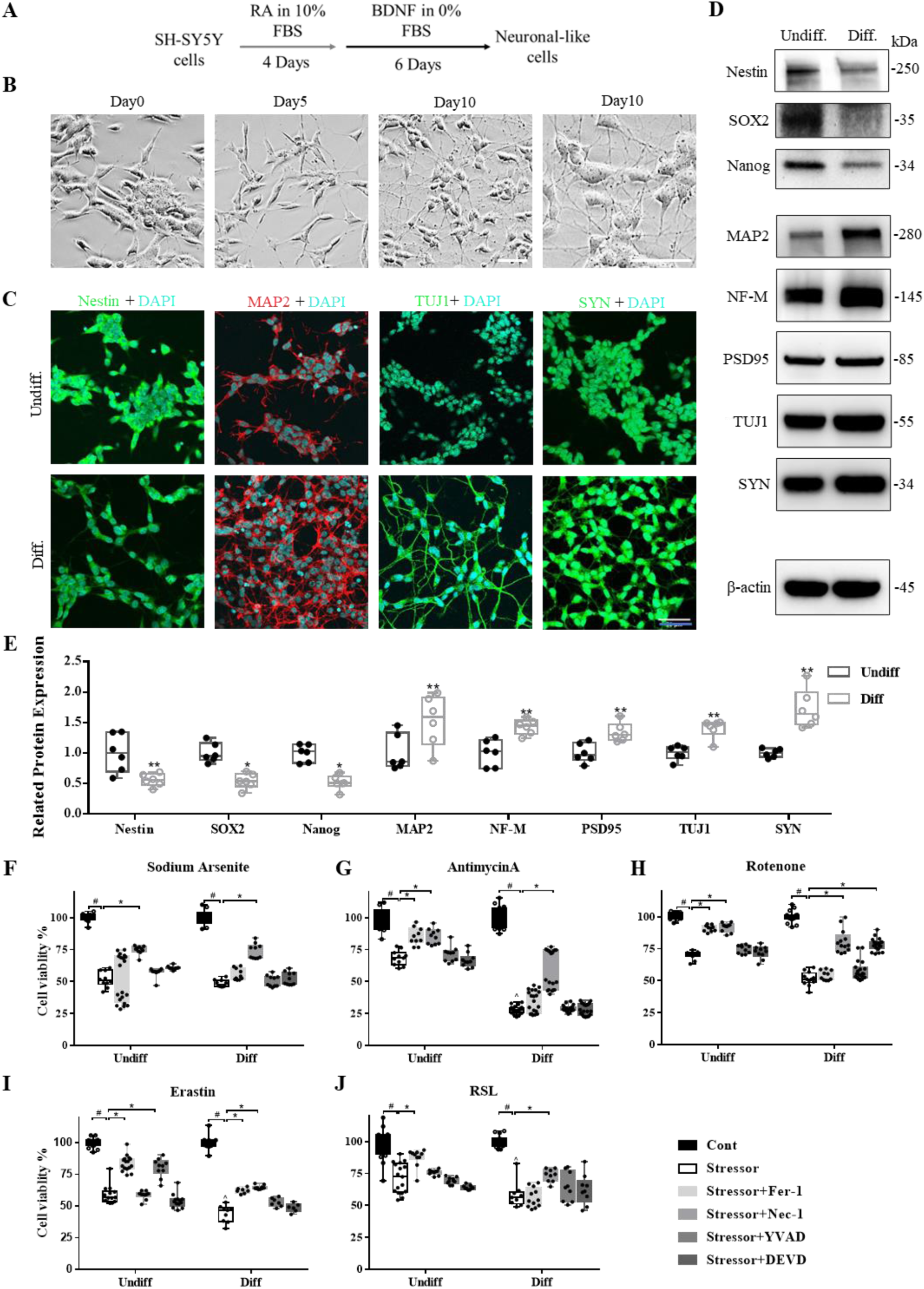
Neuronal phenotypic characterization of differentiated SH-SY5Y cells. **A** Schematic diagram of the protocol for the neuronal differentiation of SH-SY5Y cells. RA: retinoic acid; BDNF: brain derived neurotrophic factor. FBS: fetal bovine serum. **B** Morphological appearance of undifferentiated (Undiff.) (Day0) and differentiated (Diff.) (Day5, 10) SH-SY5Y cells by light microscope. Images were obtained at 20X magnification and 40X magnification. Scale bar: 50 μm. **C** Immunofluorescent staining on neuronal markers including Nestin (Green), microtubule associated protein 2 (MAP2, Red), β-tubulin-III (TUJ1, Green), Synaptophysin (SYN, Green) with DAPI (Blue) in Undiff. vs Diff. cells at Day10. Images were obtained at 40 × magnification. Scale bar: 30 μm. **D** Western blot analysis showing the lower protein expression of stem cell markers (Nestin, Nanog and SOX2) and the higher protein expression of mature neuronal markers including MAP2, Neurofilament M (NF-M), postsynaptic density protein 95 (PSD-95), TUJ and SYN in the Diff. cells. β-Actin was used as a loading control. Representative results of experiment performed twice with three biological replicates. **E** Quantification of western blot assessed by Image J software. *P < 0.05; **P < 0.01. The specific pattern of stress response and cell death in Uniff. and Diff. cells faced with Sodium Arsenite, AntimycinA, Rotenone, Erastin and 1-RSL-3 were shown in the **F**, **G**, **H**, **I** and **J**, respectively (n = 6 replicated wells, 2 independent experiments). #denotes P < 0.05 compared to the corresponding control groups. *Denotes P < 0.05 compared to the corresponding stress only groups. ^ denotes P < 0.05 compared to the Undiff. stress only groups.

We then investigated whether differentiated SH-SY5Y cells were more susceptible to oxidative stress, a characteristic commonly observed in mature neurons^19^. The results showed that treatment with Sodium Arsenite (AS, nonspecific oxidative stressor, **Figure 1F**), Antimycin A (AntiA, respiratory complex III inhibitor, **Figure 1G**), Rotenone (respiratory complex I inhibitor, **Figure 1H**), Erastin (Er, ferroptosis stressor, **Figure 1I**) and RSL (ferroptosis stressor, **Figure 1J**) significantly decreased cell viability in both Undiff. and Diff group compared to their respective control groups (^#^ P < 0.05). Interestingly, Diff. cells treated with AntiA (**Figure 1G**), Rot (**Figure 1H**), Er (**Figure 1I**) and RSL (**Figure 1J**) exhibited even lower cell viability compared to Undiff. treated with the same stressors (^ P < 0.05), indicating the heightened sensitivity of differentiated SH-SY5Y cells to these stressors. However, there was no significant difference in cell viability between Diff. and Undiff. in response to AS (**Figure 1F**), suggesting that this heightened sensitivity is stress specific. Furthermore, we also examined the type of programmed cell death by comparing the groups treated with stress inhibitors to the groups treated with stressors. Except for the reversible effect of Nec-1 (Necroptosis inhibitor) on AS-induced cell death observed in both Undiff. and Diff. group (**Figure 1F**), the activation of cell death differed between the two groups for the other 4 stressors. The effect of AntiA in Undiff. could be attenuated by both Fer-1 (Ferroptosis inhibitor) and Nec-1, while in Diff. group, it could only be reversed by Nec-1 (**Figure 1G**); Similarly, the effect of Rot in Undiff. could be attenuated by Fer-1 and Nec-1, but in Diff. group could be reversed by Nec-1 and DEVD (Pyroptosis inhibitor) (**Figure 1H**). The effect of Er in Undiff. could be attenuated by Fer-1 and YVAD (Apoptosis inhibitor), whereas in Diff., it could be reversed by Fer-1 and Nec-1 (**Figure 1I**). Lastly, the effect of RSL in Undiff. could be attenuated by Fer-1, conversely, it could only be reversed by Nec-1 in Diff. group (**Figure 1J**). Overall, the differentiation of SH-SY5Y cells towards neuron led to an enhanced sensitivity to mitochondrial stress and ferroptosis and changes in programmable cell death activation in response to these stressors.

### ➢ Transcriptome-wide mRNA Stability Profiling Reveals Differential RNA Stability in changes after differentiation

Messenger RNA stability, as a critical factor in post-transcriptional regulation, may play a significant role in the complex behavior of neuron^20^. Using a simplified differential mRNA stability profiling which combines Actinomycin D (ActD) assay with RNA sequencing, we examined the relative changes of mRNA stability across the transcriptome in both undifferentiated and differentiated state. In the Diff. vs. Undiff comparison, we identified 296 stabilized genes and 170 destabilized genes (**Figure 2A**, **Dataset 1**). To further elucidate the functional enrichment pattern of differentially stabilized genes (DSGs), we performed Gene Set Enrichment Analysis (GSEA) of Biological Process (BP), Cellular Component (CC) and Molecular Function (MF) and found stabilized genes were enriched in the pathways related to the neuronal morphogenesis and neuron’s specific functions, such as CC: Neuron projection terminus, BP: Neuromuscular synaptic transmission, BP: Regulation of dopamine secretion (**Figure 2B**). On the other hand, the functional categorization of destabilized genes was exclusively associated with BP: Mitochondria electron transport, cytochrome c to oxygen, CC: mitochondrial respiratory chain complex, glutathione metabolic process and oxidative stress relevant pathways (**Figure 2B**).

**Figure 2.**
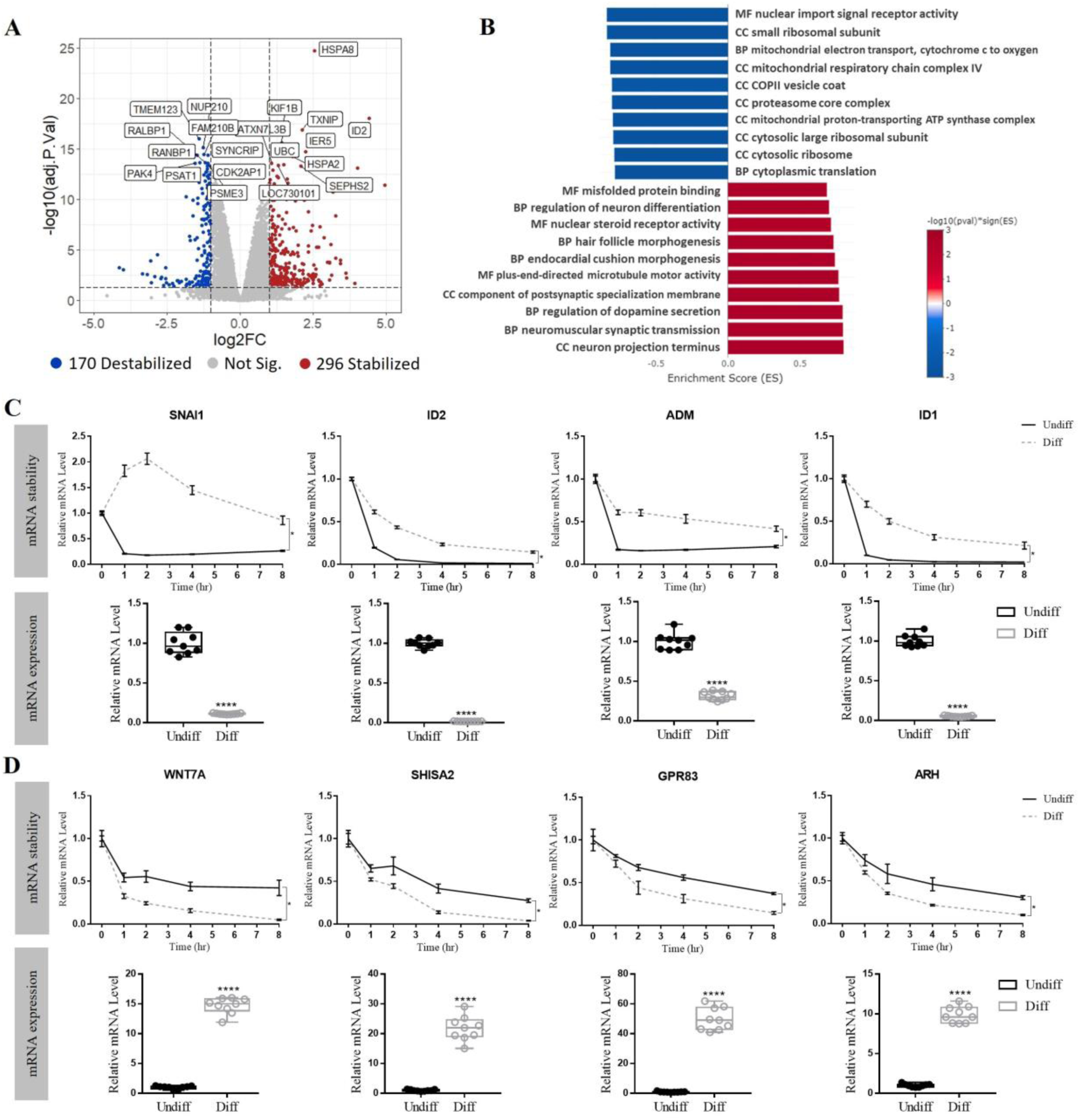
Transcriptome-wide RNA stability profiling of differentiated SH-SY5Ycells and validation. **A** The volcano plot showing the log2 FC and –log10 adj. P value of each gene. Significantly up and down differentially stabilized genes (DSGs) were indicated by red and blue dots, respectively. (Cutoff: |Log_2_FC| ≥ 1 and adj. P < 0.05). Top 10 up and down DSGs with the lowest adj. P value were labeled by gene symbol. **B** GSEA identifying Top 10 enriched GO pathways of stabilized genes (Red) and destabilized genes (Blue), respectively. **C** mRNA stability validation and gene expression of Top 4 stabilized genes in DSGs dataset by qRT-PCR. **D** mRNA stability validation and gene expression of Top 4 destabilized genes in DSGs dataset by qRT-PCR. *Denotes P < 0.05 compared to the undifferentiated group at the indicated time point. **** denotes P < 0.0001 compared to the undifferentiated group. Each condition has at least 3 biological replicates and 2 technical replicates.

Since this simplified method of transcriptome-wide mRNA stability profiling is new, the conventional “gold standard” ActD qPCR assay was employed for validation^21^. The ActD assay was conducted on the Top 4 stabilized genes and Top 4 destabilized genes (**Dataset 1**). The relative mRNA levels of SNAI1, ID2, ADM and ID1 showed a slower decrease in the Diff. group compared to Undiff. group, indicating that they were more stabilized in the Diff. group (**Figure 2C**, Upper Panel). Conversely, the relative mRNA levels of WNT7A, SHISA2, GPR83 and ARH decreased faster in the Diff. group compared to the Undiff. group, implying the destabilization of these genes in the Diff. group (**Figure 2D**, Upper Panel). These data confirmed the accuracy of this newly simplified mRNA stability profiling. Surprisingly, the steady mRNA levels of each Top 4 stabilized genes were significantly decreased in the Diff. group (**Figure 2C**, Lower Panel), while the mRNA levels of each Top4 destabilized genes were substantially increased in the Diff. group (**Figure 2D**, Lower Panel), which is counterintuitive to the known relation between mRNA half-life and mRNA expression^22^.

### ➢ mRNA Stability buffers mRNA transcription during differentiation

To better understand the position of mRNA stability in the complex regulatory network of neuronal differentiation, we conducted an integrative omics analysis of mRNA stability, transcriptomics and translatomics. Transcriptomic and translatomic alternations were detected by RNA-seq and ribosome profiling respectively and identified 1541 Differentially Expressed Genes (DEGs, Up-regulated: 992; Down-regulated:549) (**Dataset 2**, **Supplementary** Figure 1A**&B**) and 1642 Differentially Translated Genes (DTGs, Up-regulated: 946; Down-regulated: 696) (**Dataset 3**, **Supplementary** Figure 1C**&D**). Correlation analysis of all transcripts showed that mRNA stability dataset showed no significant correlated to the transcriptomic dataset and translatomic dataset (correlation coefficient: –0.053; P < 0.001) (**Figure 3A**). However, when focusing on the significantly changed genes in the three datasets (Cutoff: |log2FC| >= 1 and adj. P < 0.05), a stronger negative correlation was observed between DSGs and DEGs (correlation coefficient: –0.684; adj. P < 0.001) and between DSGs and DTGs (correlation coefficient: –0.614; adj. P < 0.001) (**Figure 3B&C**). Transcription and translation were highly correlated (correlation coefficient: 0.909; adj. P < 0.001; **Supplementary** Figure 1 **E&F**). This result indicates that RNA under-transcription and translation is counteracted by RNA stabilization, and reciprocally, RNA over-transcription and translation is counteracted by RNA destabilization when major genetic alterations were induced by neuronal differentiation, implicating the buffering effect of mRNA stability in the context of neuronal differentiation^23^.

**Figure 3.**
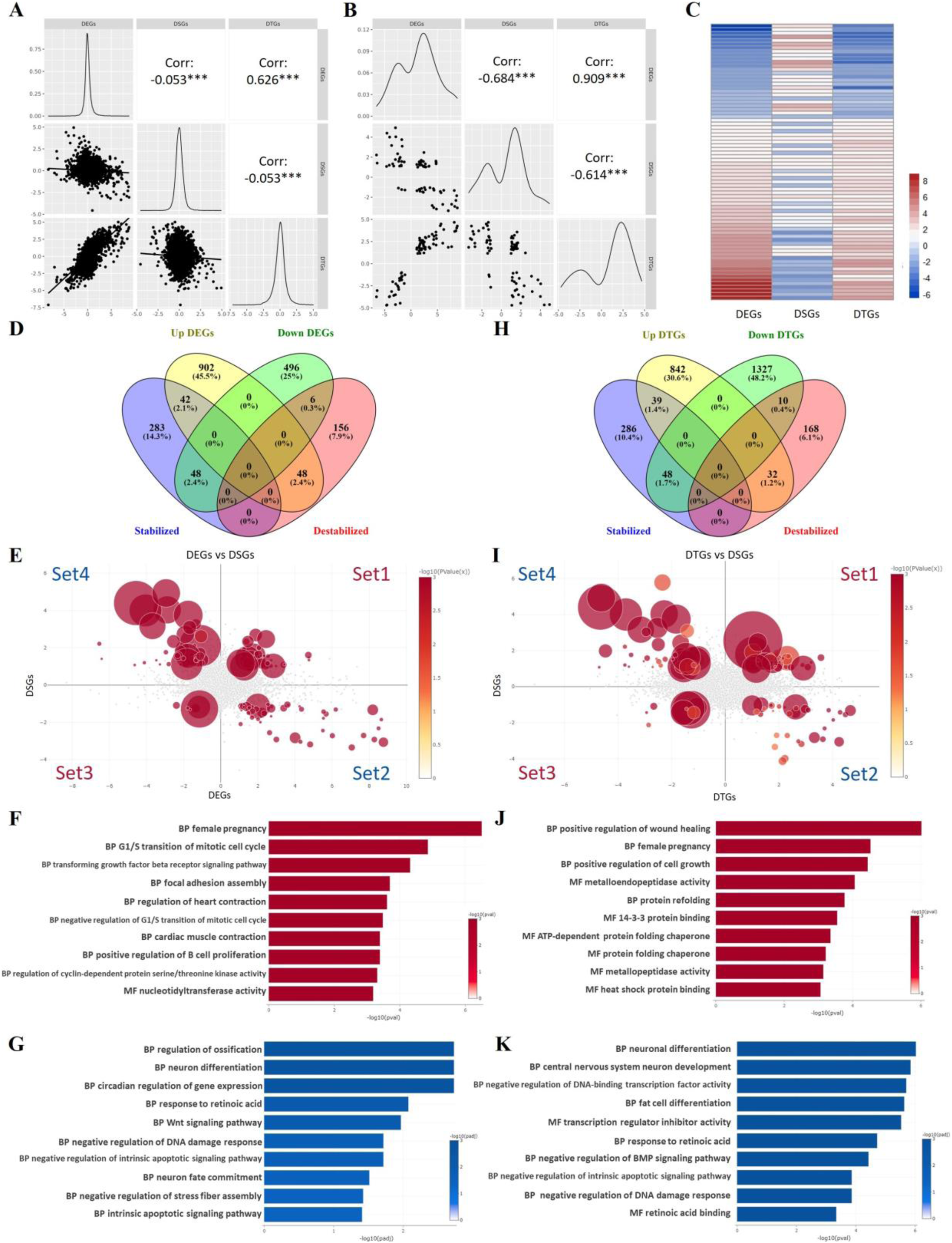
mRNA Stability Negatively Correlates with Transcriptomics and Translatomics in the Context of Neuronal Differentiation. **A** Correlation analysis of mRNA stability and transcriptome and translatome involving all the transcripts. **B** Correlation analysis among DEGs and DSGs and DTGs with applying cutoff: |Log_2_FC| ≥ 1 and adj. P < 0.05). **C** Heatmap showing Log_2_FC of DEGs, DSGs and DTGs. Venn diagrams display the ratio of overlap in DSGs vs DEGs (**D**) and DSGs vs DTGs (**H**). **E**&**I** Visualization of the Log_2_FC values of each overlapping gene between DSGs and DEGs/DTGs. The numbers indicate the territories of four sub-groups. Positive correlated groups included 1: Up-DEGs/DTGs and stabilized genes; 3: Down-DEGs/DTGs and destabilized genes. Negative correlated groups included 2: Up-DEGs/DTGs and destabilized genes; 4: Down-DEGs/DTGs and stabilized genes. **E**&**J** GO terms enrichment analysis of the positive correlated groups. The top 10 pathways were selected and displayed. **G**&**K** GO terms enrichment analysis of the negative correlated groups. The top 10 pathways were selected and displayed.

We conducted a more detailed analysis of the relationship between DSGs vs DEGs, as well as DSGs vs DTGs. We found that there were 110 genes that were shared between DSGs and DEGs, and out of these, 76 genes showed a significant change in an opposite direction (**Figure 3D**). We then visualized these genes into the two-dimensional space of DSGs and DEGs and categorized them into two groups based on their correlation with DEGs. The Positive Correlation Group consisted of two sets: Set1: Up-DEGs and Stabilized Genes (29 genes, 1.5%) and Set3: Down-DEGs and Destabilized Genes (5 genes, 0.3%). On the other hand, the Negative Correlation Group consisted of Set2: Up-DEGs and Destabilized Genes (39 genes, 2.1%) and Set4: Down-DEGs and Stabilized Genes (37 genes, 2%) (**Figure 3D&E**). To further investigate the functional implications of these gene groups, we conducted an Over-Representation Analysis (ORA) on the genes belonging to different sets. The Positive Correlation Group was mainly associated with cell proliferation and cell cycle related pathways (**Figure 3F**), while Negative Correlation Group clustered in neuronal differentiation-related pathways and stress-related pathways (**Figure 3G**). This result suggests the possible bidirectional regulatory mechanism of mRNA stability on transcriptomics, and the explicit direction of regulation is gene function and physiological condition dependent. Given the highly positive correlation between DTGs and DEGs, the analysis of DSGs vs DTGs showed that Positive Correlation Group was formed by 49 genes and Negative Correlation Group contained 80 genes which were consistently enriched in the cell growth-related pathways and neuronal differentiation-associated pathways (**Figure 3H-K**). This data points out the processes of mRNA decay and translation are potentially coupled in the context of neuronal differentiation.

The positive correlations of mRNA stability with transcripts abundance and the process of translation have been observed in various organisms under the different states, while the counteracting effect of mRNA stability is out of expectation, it was nonetheless previously reported, but without enough validation^24^. Hence, based on the functional categorization of Negative Correlation Group, we selected WNT7A (representative of neuronal differentiation-related genes) and SEPHS2 (representative of neuronal stress-related genes) as the main targets to verify the observations acquired from the bioinformatic analysis. WNT7A was the Top 1 significantly destabilized gene, reported to play a crucial role in mediating neuronal differentiation ^25^. The mRNA and protein expression levels of WNT7A were significantly increased in Diff. vs Undiff. group, while the stability of WNT7A changed in the opposite direction (**Figure 4A**).

**Figure 4.**
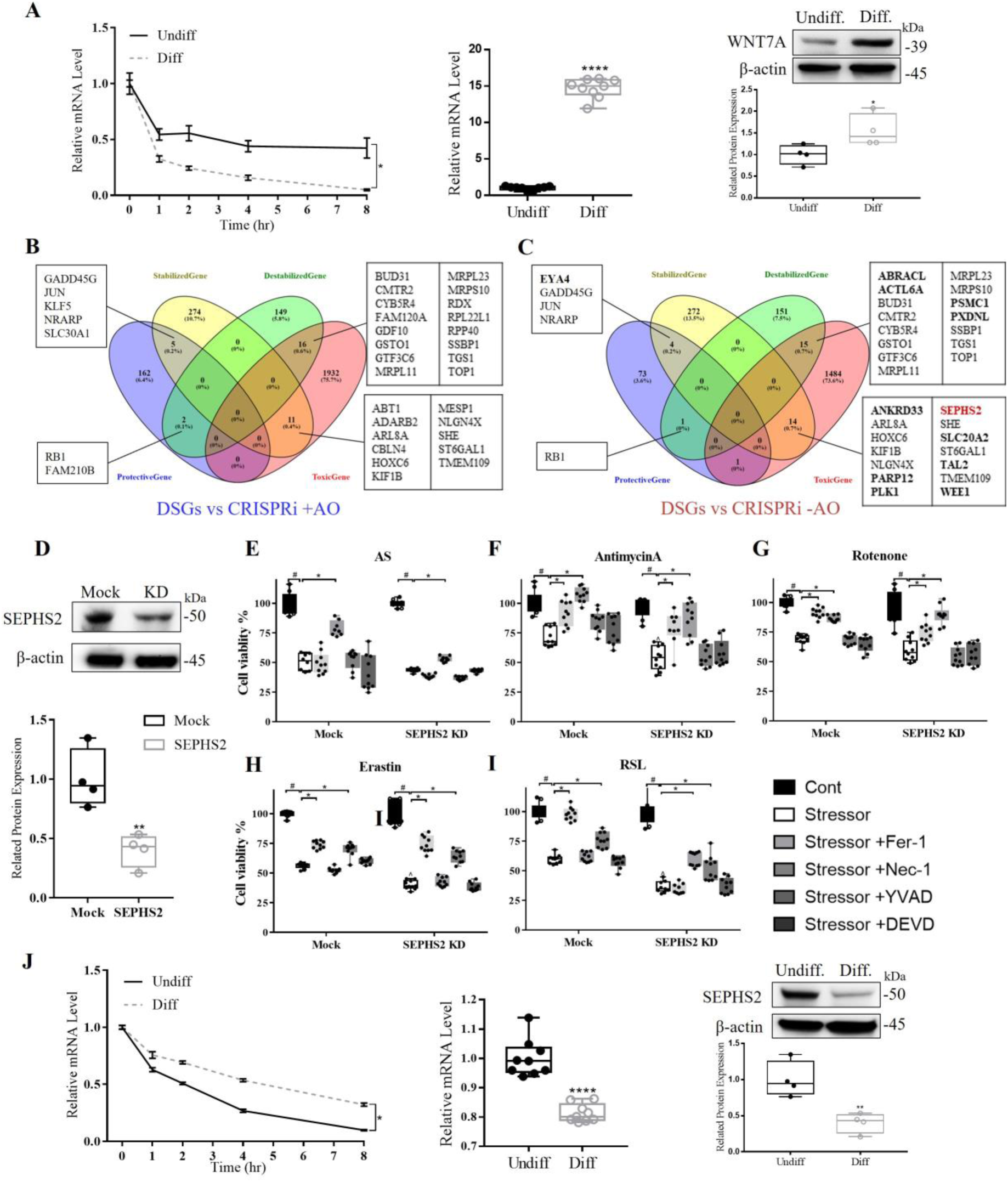
WNT7A and SEPHS2 Validates the Buffering Effect of mRNA stability on Gene Expression and Protein Expression. **A** The gene expression, protein expression and mRNA stability of WNT7A showing around 15-fold increase of mRNA and 1.5-fold increase of protein while destabilization of WNT7A. **B** Venn diagrams displaying the ratio of overlap between DSGs in differentiated SH-SY5Y cells and survival genes of CRISPR inference plus antioxidants in iPSCs-derived neurons. **C** Venn diagrams display the ratio of overlap between DSGs in differentiated SH-SY5Y cells and survival genes of CRISPR inference minus antioxidants in iPSCs-derived neurons. **D** Knockdown of SEPHS2 (KD) protein expression by infecting shRNAs/lentivirus and mock-shRNA/lentivirus in SH-SY5Y cells. Knockdown efficiency was quantified based on the band intensity by ImageJ. SEPHS2 KD showing the enhanced neuronal vulnerability to stress and distinct cell death program. The specific stress response and cell death type of Mock and SEPHS2 treated with Sodium Arsenite, Antimycin A, Rotenone, Erastin and 1-RSL-3 were shown in the **E**, **F**, **G**, **H** and **I**, respectively. Cell viability was expressed as mean ± SD (n = 6 replicated wells, 2 independent experiments). # denotes P < 0.05 compared to the corresponding control groups. * Denotes P < 0.05 compared to the corresponding stress only groups. ^ denotes P < 0.05 compared to the Mock stress only group. **J** The gene expression, protein expression and mRNA stability of SEPHS2 displaying 1.2-fold decrease of mRNA and 2-fold decrease of protein while stabilization of SEPHS2, suggested the buffering role of mRNA stability between transcription and translation.

On the other hand, to accurately screen out the crucial neuronal stress-related genes, and identify the potential of mRNA dynamic stability changes in regulating this process, we further analyzed the mRNA stability dataset against a published genome-wide CRISPR screening dataset on iPSC-derived neurons^26^. CRISPR inference (CRISPRi) enabled a large-scale loss-of-function genetic screens and uncovered numerous genes controlling neuronal response to chronic oxidative stress. Overlapped analysis of DSGs vs CRISPRi plus antioxidants (+AO) and DSGs vs CRISPRi no antioxidants (-AO) were performed individually. The overlapped genes in DSGs vs CRISPRi +AO showed a high similarity to those in DSGs vs CRISPRi –AO (**Figure 4B&C**). We then focused on the 12 genes which were unique to DSGs vs CRISPRi – AO as the latter one was established in a more physiologically relevant approximation of chronic oxidative stress. Among these 12 genes, SEPHS2 was the most significant gene, with little direct evidence that SEPHS2 could affect neuronal survival under oxidative stress^27^. Next, we identified that knockdown of SEPHS2 sensitized the undifferentiated cells to mitochondrial stress and ferroptosis (**Figure 4D-I**). Accordingly, the mRNA and protein expression level of SEPHS2 were predominantly decreased in the Diff. group compared to Undiff. group, while it was more stabilized after the differentiation (**Figure 4J**). These results further confirmed that the dynamic changes in mRNA stability may counteract the alterations of transcription on translation which were induced by neuronal differentiation, and also demonstrate the value of mRNA stability in uncovering important regulators of the studied phenotypes.

### ➢ RNA Binding Proteins regulate the dynamic mRNA Stability changes

In mammalian cells, mRNA stability largely depends on the mRNA nucleotide sequence, which determines the codon compositions and impacts the accessibility of RNA-binding proteins to the mRNA^28^. Given that the coupling of mRNA half-life to translation is mediated by codon optimality of mRNAs^29^, we investigated the codon usage and optimality of differentially stabilized/destabilized mRNAs by analyzing synonymous codon usage and global codon usage. It was clearly observed that some codons preferentially occurred in differentially stabilized mRNAs while others were more common in differentially destabilized mRNAs (**Supplementary** Figure 2A). Through heatmap plots and hierarchical clustering, we observed distinct patterns of codon over-or under-usage in each dataset of stability, translation (Ribo-seq) and translational efficiency (TE). Clustering analysis of codon usage also revealed a negative correlation of Stability Up-Ribo Up and Stability Up-TE Up, as well as a positive correlation of Stability Down-Ribo Down and Stability Down-TE Down) (**Supplementary** Figure 2B**&C**). Moreover, the global codon usage patterns showed a similar phenomenon as the synonymous codon usage analysis (**Supplementary** Figure 3D**&E**). While we observed that DSGs are codon biased, it is clear that this bias does not correlate with the apparent codon biases and optimality observed at the level of mRNA translation, indicating that the impact of dynamic mRNA stability changes on mRNA translation is not driven by codon usage patterns, or at least is not a main contributor to such relation.

In recent years, RBPs have been increasingly recognized for their pivotal role in interacting with specific mRNA sequences and therefore regulating mRNA stability machinery^30^. We performed the motif enrichment analysis on the mRNA stability dataset to explore the potential RBPs that may influence the neuronal differentiation outcome and stress response^31^. Changes in the transcript stability were ranked based on the signal-to-noise ratio, where transcripts stabilized in the Diff. cells had positive values and those destabilized had negative values. k-mer-based Transcript Set Motif Analysis (TSMA) of differentially stabilized genes, revealed a set of enriched k-mers in the differentiated cells that were largely CG-rich (**Supplementary Table 1**). These k-mers mapped to the motifs of SAMD4 and SRSF as the Top 2 hits (**Figure 5A, Supplementary** Figure 3). Furthermore, the k-mers corresponding to the SAMD4-binding motif, shown in yellow, were among the most highly enriched k-mers in the transcripts that were found to be stabilized (**Figure 5B**). This was validated by Spectrum Motif Analysis (SPMA), which revealed these same Top 2 RBPs (**Figure 5C, Supplementary** Figure 4). The spectrum plot of Top 1 RBP SAMD4A/B revealed a highly consistent, nearly monotonic increase in SAMD4-binding sites when the genes were ranked from those most destabilized to those most stabilized after SH-SY5Y differentiation (**Figure 5D**). Specifically, the putative binding sites of SAMD4 were highly enriched in stabilized transcripts (shown in red) and highly depleted in destabilized transcripts (shown in blue) in the differentiation cells. Based on these results, SAMD4A/B emerged as the single best RBP out of the 174 RBPs in the database whose motif could rationalize the observed changes in stability.

**Figure 5.**
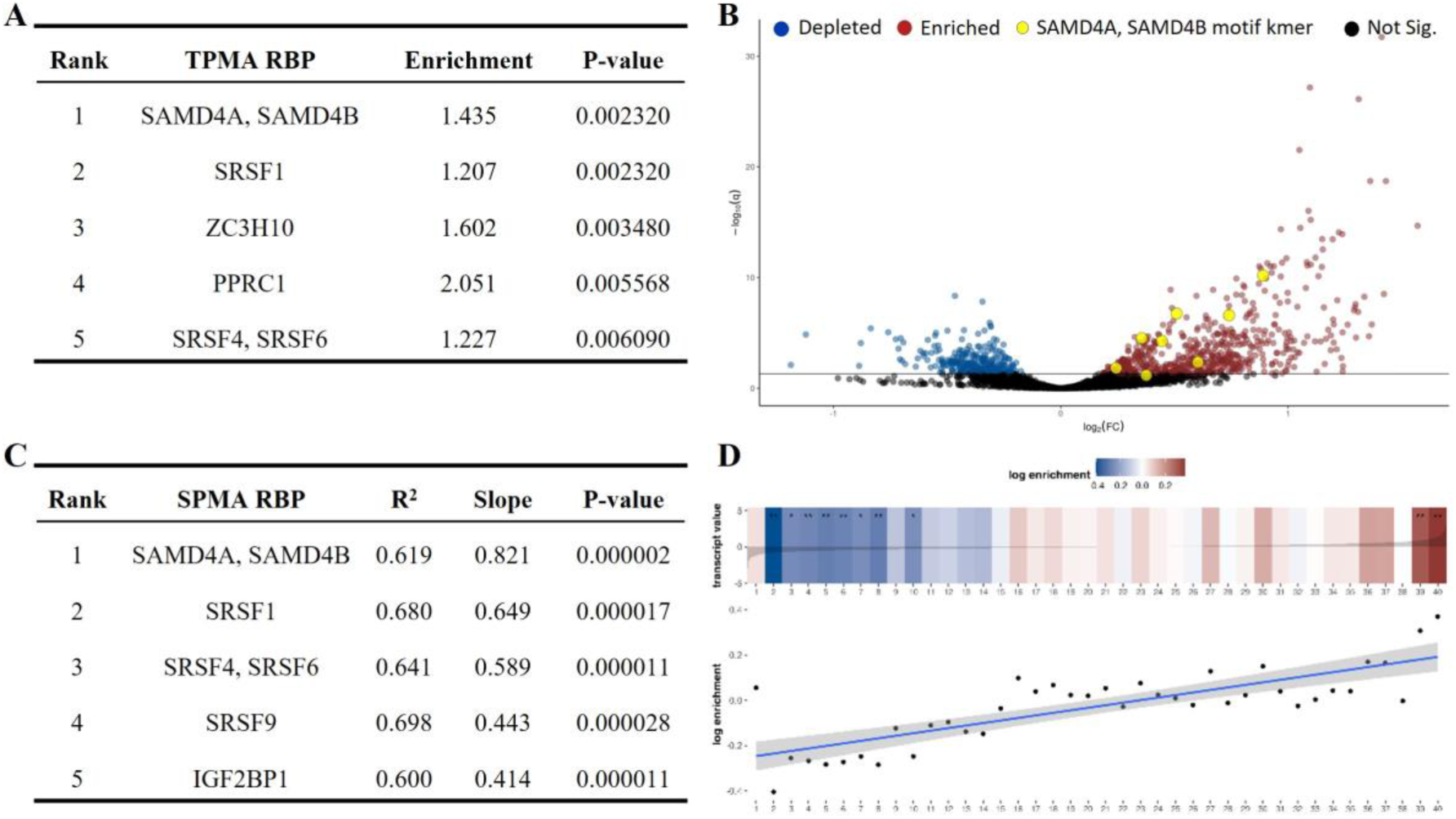
Spectrum Motif Analysis (SPMA) identified SAMD4A motifs as highly enriched in stabilized gene sets in the differentiated cells. **A** displaying Top5 RBPs with highly enriched motifs in the stabilized genes for Transcript Set Motif Analysis (TSMA). **B** TSMA volcano plot showing enriched and depleted k-mers in stabilized genes after the differentiation. k-mers associated with SAMD4A, SAMD4B (shown in yellow are highly enriched). **C** displaying Top5 RBPs with highly non-random motif enrichment patterns for SPMA. **D** SPMA depicting the distribution of putative binding sites of SAMD4A across all the transcripts. The transcripts are sorted by ascending signal to noise ratios. The transcripts destabilized in Diff. groups relative to Undiff. group are on the left, and those stabilized are on the right of the spectrum. The putative binding sites of SAMD4A are highly enriched in transcripts stabilized in Diff group (shown in red) and highly depleted in transcripts destabilized in Diff group (shown in blue).

### ➢ SAMD4A regulates neuronal differentiation and response to oxidative stress

Due to the lack of direct evidence on the role of SAMD4A in neuronal differentiation, we investigated the differentiation outcome and stress response after knocking down SAMD4A to test the validity of our approach. We observed that SAMD4A expression significantly increased following the SH-SY5Y differentiation process (**Figure 6A**). Using short hairpin RNA (shRNA), we successfully silenced SAMD4A expression by 66% in undifferentiated SH-SY5Y cells (**Figure 6B**). As a result, we observed a dramatic increase in the expression of stem cell markers (Nestin, Nanog and SOX2), but no significant changes in mature neuronal markers (MAP2, NF-M, PSD95, TUJ1 and SYN) in the SAMD4A knockdown group (SAMD4A KD) compared to the Mock group (**Figure 6C, Supplementary** Figure 5). With regard to the morphological alternations, there were no obvious differences between SAMD4A KD and its Mock group before induction of differentiation. However, during the RA stage of pre-differentiation, SAMD4A KD cells started to exhibit significantly fewer neurites compared to the Mock group (**Figure 6D**). Treatment with BDNF exacerbated the differences between SAMD4A KD and Mock group, as demonstrated by the impaired neurite outgrowth and sparse neurite network (**Figure 6D**). Additionally, quantitative analysis of neurite outgrowth consistently showed a reduced growth rate throughout the whole differentiation process in the KD cells (**Figure 6E**). Time course analysis during differentiation revealed that Nestin, Nanog and SOX2 continued to be highly expressed, while MAP2 and SYN were consistently expressed at a low level along the process of differentiation in SAMD4A KD cells (**Figure 6F**). This suggests that cells were unable to overcome the stemness induced by SAMD4A KD thus failed to initiate the normal differentiation. In conclusion, knocking down SAMD4A resulted in the persistent maintenance of stemness and dysregulation of neuronal differentiation.

**Figure 6.**
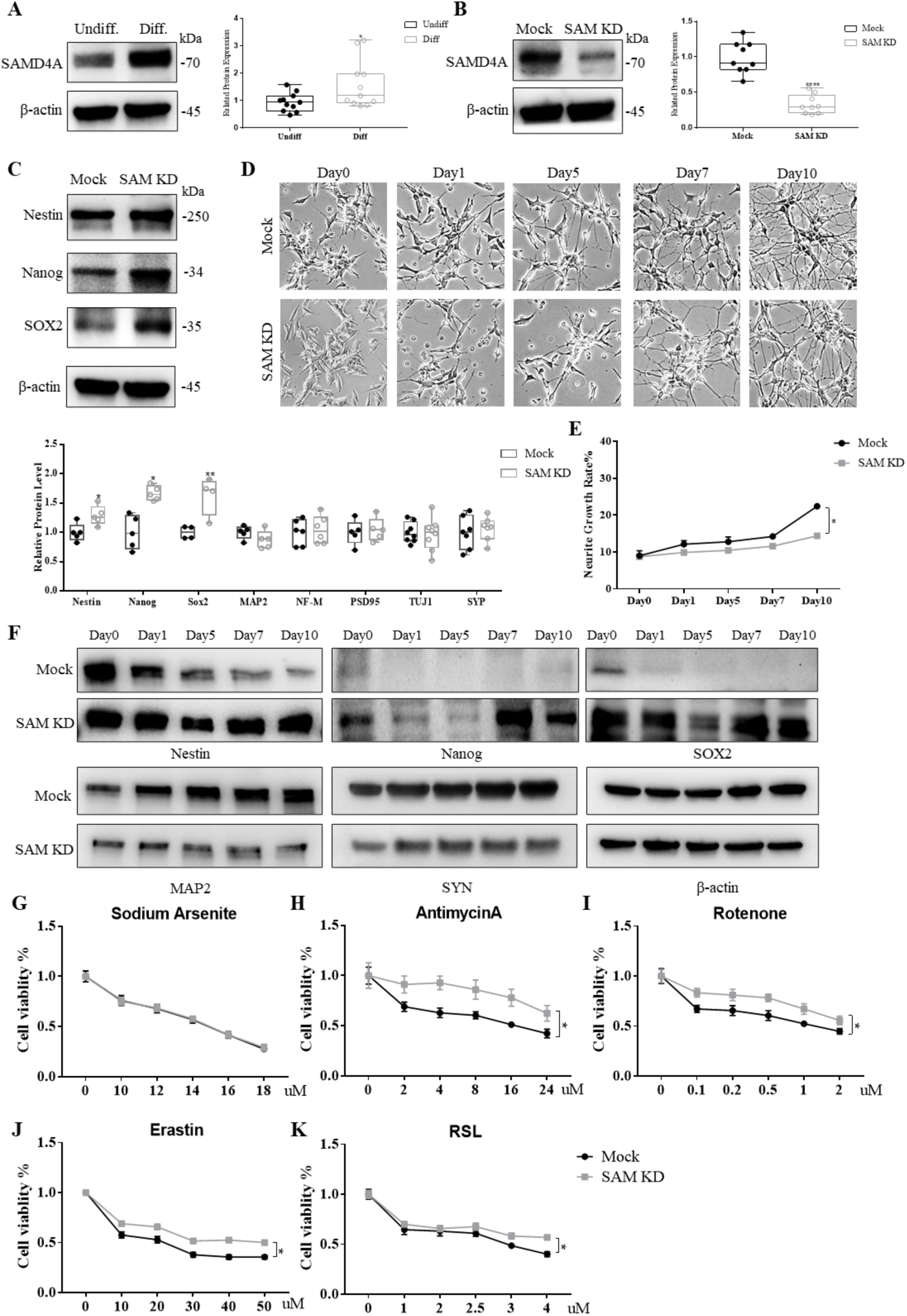
SAMD4A Knockdown (KD) Contributes to Maintained Stemness and Impaired Neuronal Differentiation and Renders Resistance to Oxidative Stress. **A** Western blot analysis showing SAMD4A expressed higher after the neuronal differentiation. Quantification was assessed by ImageJ. **B** Knockdown of SAMD4A protein expression by infecting SAMD4A-shRNAs/lentivirus and mock-shRNA/lentivirus in SH-SY5Y cells. Knockdown efficiency was quantified based on the band intensity by ImageJ. **C** The protein expression of Nestin, Nanog and SOX2 was measured in the SAMD4A KD and Mock cells, respectively. Quantification of western blot showing the up-regulated protein expression of Nestin, Nanog and SOX2 after SAMD4A KD, while no difference of mature neuronal markers MAP2, NF-M, PSD95, TUJ1 and SYN. **D** The morphologic alternations of SAMD4A KD and Mock cells during the differentiation showing the sparse neurite outgrowth. **E** Quantification of neurite growth rate based on the **D** showing the decrease neurite growth rate in the SAMD4A KD group. **F** Time course analysis of expression of Nestin, Nanog, SOX2, MAP2 and SYN during the whole process of neuronal differentiation. **G-K** The effect of SAMD4A KD on the stress response. No difference of the cell viability was observed between SAMD4A KD and Mock group in the response to sodium Arsenite (**G**). **H**, **I**, **J** and **K** represented that SAMD4A KD rendered cells more resistant to the stress induced by Antimycin A, Rotenone, Erastin and RSL3. Cell viability was expressed as mean ± SD (n = 6 replicated wells, 2 independent experiments).

Furthermore, we observed the effect of SAMD4A on the neuronal sensitivity to oxidative stress. In response to AS, both SAMD4A KD and Mock groups showed a significant decrease in cell viability as the concentration of AS increased, with no difference in cell viability between these two groups (**Figure 6G**). On the other hand, when exposed to stress induced by AntiA (**Figure 6H**), Rot (**Figure 6I**), Er (**Figure 6J**) and RSL (**Figure 6K**), SAMD4A KD cells exhibited higher cell viability. This suggests that SAMD4A may play a crucial role in influencing neuronal sensitivity to oxidative stress either directly or indirectly by regulating the differentiation phenotype.

### ➢ SAMD4A fine-tunes mRNA stability to regulate neuronal function and differentiation

Given the observed connection between SAMD4A and neuronal behavior, we further investigated the underlying mechanism by RNA sequencing and RNA stability profiling after SAMD4A KD. Differential gene expression analysis identified 117 genes (33 upregulated, 84 downregulated) in SAMD4A KD versus Mock cells (|Log_2_FC| ≥ 1.5 and adj. P < 0.05, **Figure 7A, Dataset 4**). GSEA of upregulated DEGs showed an enrichment for functions related to cell cycle pathways, while downregulated DEGs functionally enriched in the neuronal morphogenesis-associated pathways (regulation of neuronal synaptic plasticity) and more importantly, oxidative phosphorylation-related (mitochondrial electron transport) and programmed cell death-related pathways (intrinsic apoptotic signaling and P53 signaling pathways) (**Figure 7B**). This data reveals key molecular events underlying the effect of SAMD4A on promoting neuronal differentiation and sensitizing neurons to stress. On the other hand, through global mRNA stability profiling: 53 stabilized and 45 destabilized mRNAs were revealed in SAMD4A KD cells (|Log_2_FC| ≥ 1.5 and adj. P < 0.05, **Figure 7C, Dataset 5**). GSEA of stabilized genes indicated that many of the top enriched gene ontology (GO) terms related to cellular components and biological processes including mitochondrial respiratory chain and neuron apoptotic process (**Figure 7D**). Destabilized genes corresponded to processes involved in cell proliferation, such as cell cycle and cell division (**Figure 7D**). The enrichment patterns of Stabilized DSGs and Destabilized DSGs in SAMD4A KD cells were exactly opposite to those of DSGs in differentiated cells, where SAMD4A was highly expressed. This phenomenon demonstrates the impact of SAMD4A on mRNA stability in the context of neuronal differentiation.

**Figure 7.**
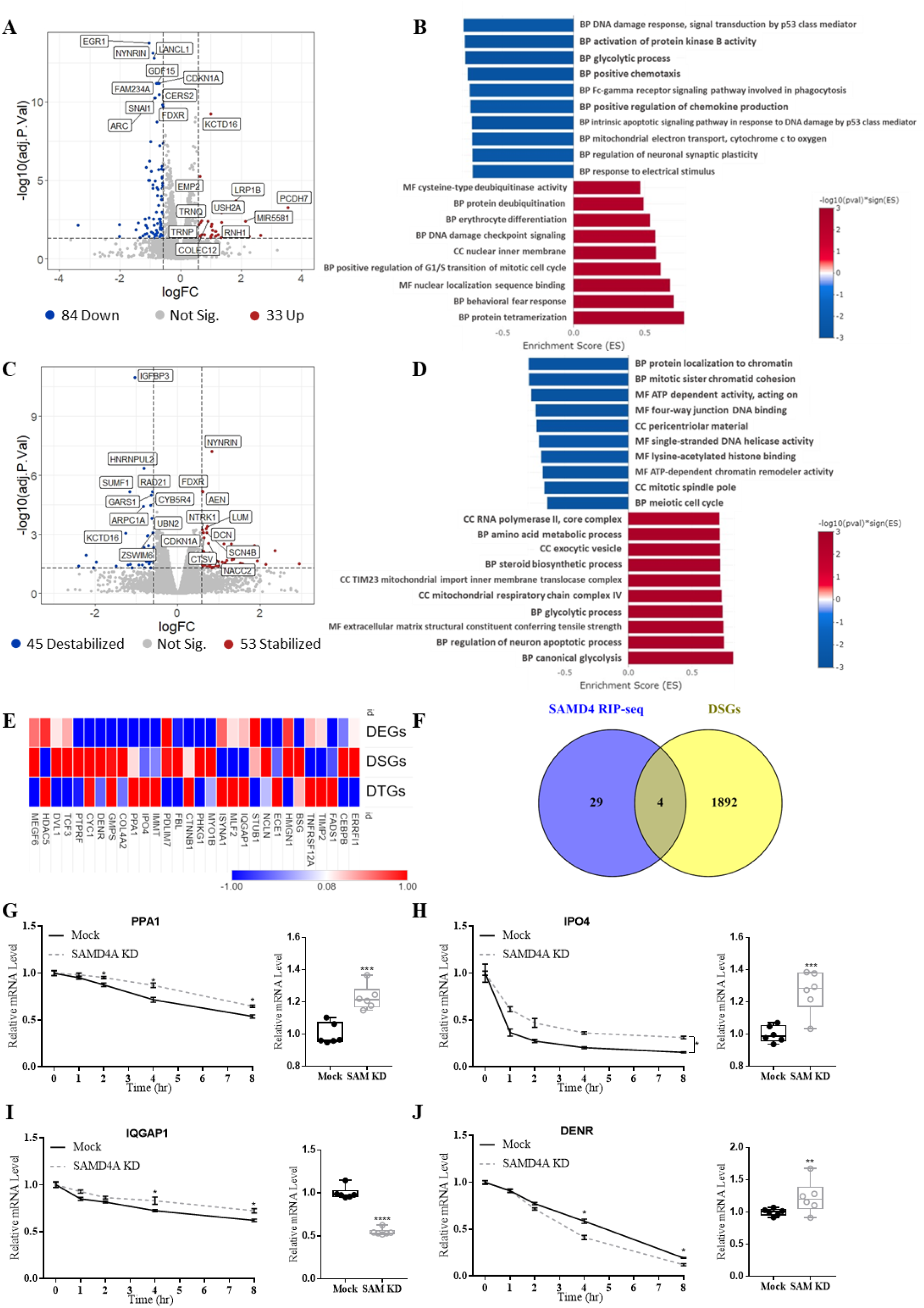
SAMD4A Regulates Neuron’s Behaviors by Tuning mRNA stability. **A** The volcano plot showing the log2 FC and –log10 adj. P value of DEGs in SAMD4A KD vs Mock. (Cutoff: |Log_2_FC| ≥ 1 and adj. P < 0.05). Top 10 up and down DEGs with the lowest adj. P value were labeled by gene symbol. **B** GSEA identifying Top 10 enriched GO pathways of Up-DEGs (Red) and Down-DEGs (Blue), respectively. **C** The volcano plot showing the log2 FC and –log10 adj. P value of DSGs in SAMD4A KD vs Mock. (Cutoff: |Log_2_FC| ≥ 1 and adj. P < 0.05). Top 10 up and down DSGs with the lowest adj. P value were labeled by gene symbol. **D** GSEA identifying Top 10 enriched GO pathways of Stabilized genes (Red) and Destabilized genes (Blue), respectively. **E** Heatmap of Log_2_FC of RIP-seq genes in the dataset of transcriptome, stability and translatome. **F** Venn diagram showing 4 overlapped genes between the published RNA Immunoprecipitation sequencing (RIP-seq) and DSGs datasets. **G**, **H, I** and **J** representing the mRNA expression and stability of PPA1, IPO4, IQGAP1 and DENR. SAMD4A KD could stabilize PPA1, IPO4 and IQGAP1 while destabilize DENR. The relationship between mRNA stability and gene expression followed the different two patterns: negative correlation (IQGAP1 and DENR) and positive correlation (PPA1 and IPO4).

Next, to further validate our findings, we analyzed a previously published RNA-SAMD4A Immunoprecipitation Sequencing (RIP-seq) dataset^32^. In this dataset, 33 SAMD4A-bound RNAs were identified (**Dataset 6**). Heatmap showing Log_2_FC of these SAMD4A bound genes in the Wild-Type differentiated vs undifferentiated datasets revealed the negative correlation between stability of SAMD4A bound mRNAs and their transcriptional levels as well as the buffering effect of stability changes on translation levels (**Figure 7E**). This was further evident by the correlation analysis of DEGs and DSGs after SAMD4A KD (correlation coefficient of all genes: –0.581; correlation coefficient of differential genes: –0.878; adj. P < 0.001; **Supplementary** Figure 6). The analysis of the RIP-seq and DSGs from SAMD4A KD cells (Cutoff: |Log_2_FC| ≥ 0.5 and adj. P < 0.01) datasets revealed that 4 genes (PPA1, IPO4 DENR, and IQGAP1) were common to both datasets (**Figure 7F**), whose stability were not previously associated with neurogenesis and survival. It is now a question of whether SAMD4A could influence the stability and expression of these target genes. The stability of PPA1, IPO4, and IQGAP1 was significantly increased in SAMD4A KD cell (**Figure 7G&H&I**), while DENR became more destabilized after SAMD4A KD (**Figure 7J**. This implies that SAMD4A acts as a stabilizer to DENR and a destabilizer to PPA1, IPO4 and IQGAP1. Additionally, the mRNA expression levels of these genes showed two opposite patterns in relation to mRNA stability. IQGAP1 with lower expression and greater stability in SAMD4A KD cells and DENR with higher expression and less stability in SAMD4A KD displayed the negative correlation between gene expression and RNA stability; Conversely, PPA1 and IPO4 exhibited a positive correlation between mRNA expression and stability, with higher expression mad more stability in SAMD4A KD cells. These findings suggest that SAMD4A may influence the expression of its specific genes via bi-directionally modulating their mRNA stability and therefore play a vital role in promoting neuronal differentiation and sensitizes neurons to oxidative stress.

## Discussion

In this work, we addressed the essential role of mRNA stability during neuronal differentiation and how understanding this process elucidated post-transcriptional regulators of increased susceptibility of neurons to oxidative stress. Our newly simplified mRNA differential stability profiling allows the measurement of altered RNA stability that happens at the system level in dynamic and changing conditions with few experimental resources. More interestingly, we observed negative correlation between RNA stability and both transcriptome and translatome in the context of human neuronal differentiation. While this was reported before in prokaryotes^24^, it was not studied to that extent in complex systems as the one used herein, nor was it validated or its regulators identified when reported in eukaryotes^23^. This provides an insight into understanding the regulatory role of dynamic mRNA stability changes. That is mRNA stability may function as a buffering system between transcription and translation to regulate gene expression. RBPs may be key upstream regulators to modulate mRNA stability in neuronal differentiation and other systems to ensure that the translational machinery is not over or underwhelmed by drastic changes in mRNA transcription, thus maintaining the cellular functional proteome.

### The Newly Simplified transcriptome-wide mRNA stability approach focuses on the differential stability alterations

With recent advancements in high-throughput sequencing, our ability to comprehend mRNA stability across the transcriptome has significantly improved. There are two commonly used transcriptome-wide mRNA stability analysis techniques: metabolic labeling using nucleoside analogues and transcription inhibition using the Actinomycin D^33^. However, it is important to acknowledge the drawbacks of these approaches. First, they involve the exogenous RNA sequencings (RNA spike-ins) into the library for the normalization. Second, they allow for the calculation of absolute RNA half-lives, but they are limited to comparing differential mRNA stability between different conditions, which requires multiple analysis steps to deduce. Third, the previously used methods are resources heavy. For example, in the Actinomycin D-based method, several time points are used, each with duplicate or triplicate samples, leading to exponential increase in sequencing costs and hands-on time. In this study, we optimized the approach for measuring transcriptome-wide differential mRNA stability, which was modified from the previous transcription shutoff methods. The algorithm involves a two-step normalization process. In the first step, the Actinomycin D treated libraries are normalized to the untreated libraries, Next, the two conditions are compared to identify DSGs. This allows for the analysis of differential mRNA stability in dynamic systems, in this case we compared the undifferentiated and differentiated states. Although our approach is difficult to calculate the exact mRNA decay rates (i.e., mRNA half-lives), it still greatly simplifies the existing mRNA stability profiling approach and provides valuable data for understanding the dynamic nature of mRNA stability, allowing its seamless incorporation into experimental workflow, thus providing a powerful tool to understand the complex nature of gene expression regulation. Indeed, we conducted various validation steps to validate this simplified approach, which shows, as presented in this work, its robustness and importance in providing valuable information on post-transcriptional regulatory processes.

### ➢ Transcriptome-wide mRNA Stability Profiling Reveals Neuronal Specific mRNA Decay Kinetics

In our analysis, we observed many transcripts with increased stability are functionally associated with the GO categories of neuron projection terminus, synaptic transmission, and dopamine secretion, which aligns with the characteristics of mature neurons. Many of these genes are specifically or highly expressed in the central nervous system. Examples include *Synaptotagmin 1* (SYT1), *Discs large MAGUK scaffold protein 4* (DLG4), and *Kinesin family member 1B* (KIF1B). SYT1 gene encodes the neural specific protein participating in triggering neurotransmitter release at the synapse in response to calcium binding^34^. DLG4 is also exclusively expressed in neuronal tissues at postsynaptic sites, where it exerts its function on synaptogenesis and synaptic plasticity^35^. KIF1B is detected ubiquitously in various tissues, with high levels of protein concentrated in the neuron projections^36^. The stability of these transcripts may support appropriate levels of protein production in differentiated SH-SY5Y cells without the need for high rates of transcription and translation. On the other hand, transcripts with decreased stability are predominantly clustered in ribosome biogenesis and mRNA metabolism, which are closely linked to cell growth, proliferation, and differentiation^37,38^. This may be related to the transition from proliferation to differentiation and maturation in SH-SY5Y cells. By destabilizing mRNAs involved in transcription and translation, cells can rapidly respond to differentiation inducers and generate dynamic molecules to initiate the differentiation process. This neural specific decay kinetics has also been observed in Drosophila neural development, where mRNA stability was detected by pulse-chase approach termed “TU-decay”^13^. These findings offer a new perspective on understanding neuronal differentiation and could be further explored in the progression of neuronal lineage in stem cells.

### ➢ Altered mRNA Stability impacts the Selective neuronal Vulnerability to Oxidative Stress

Beyond the established neuronal phenotype of SH-SY5Y differentiation, one of the most interesting, certainly more important findings in this work, is the heightened sensitivity of differentiated SH-SY5Y cells in response to oxidative stress. In addition to being used as a model for neuronal differentiation, SH-SY5Y cells have also been identified as an *in-vitro* model for neurodegenerative diseases, particularly Parkinson’s disease^39^. This is due to their susceptibility to mitochondrial dysfunction and high vulnerability to oxidative stress after undergoing differentiation. However, previous studies on the vulnerability of differentiated SH-SY5Y cells have primarily focused on verifying their stress responses or exploring mechanism in low throughput approaches, resulting in limited understanding of the increased sensitivity of oxidative stress^39,40^. In this study, we systematically characterized the stress response and the programmed cell death of both undifferentiated and differentiated SH-SY5Y cells by employing five different stressors from three categories, along with four cell death inhibitors. When exposed to AS, a general oxidative stressor, both undifferentiated and differentiated cells similarly exhibited a decreased cell viability, which may be mainly caused by the same cell death type necroptosis. Apart from sodium arsenite, differentiated cells showed enhanced sensitivity to other four stressors, such as Antimycin A (inhibitor of mitochondrial electron transport chain complex III), Rotenone (inhibitor of mitochondrial electron transport chain complex I), Erastin (inhibitor of cystine/glutamate antiporter and glutathione synthesis) and RSL3 (inhibitor of the glutathione peroxidase 4). Furthermore, the dissimilar cell death programs observed in differentiated cells highlighted the specific molecular cascades involved in the neuronal cell death ^41^, further contributing to their increased vulnerability to stress.

mRNA stability was shown to play a role in bacterial and yeast responses to stresses^42,43^, however, this was not studied in higher eukaryotes to the best of our knowledge. Our mRNA stability profiling collectively unraveled exclusive destabilization of genes involved in mitochondrial metabolism and function in human neuroblastoma cells differentiation. Most interestingly, our differentially stabilized gene sets partially overlap with survival genes controlling neuronal response to chronic oxidative stress uncovered by CRISPRi screen on iPSC-derived neurons^26^. Importantly, SEPHS2, which was shown to be a gene of interest in this analysis, did not show drastic transcriptional dysregulation after differentiation, thus utilizing RNA-seq data did not reveal such links. This is a testament to the importance of understanding the post-transcriptional process of mRNA stability regulating in understanding neuronal behavior and in understanding disease pathophysiology. Understanding how RNA stability regulation contributes to the vulnerability of neurons to adverse stimuli is an emerging question^44^. In physiological states, RNA homeostasis is the outcome of the intricate balance between stability promoting factors and decay factors, both of which are controlled by RNA binding proteins (RBPs)^45^. This balance is of particular importance in neurons, which are among the most metabolically active and morphologically complex cells. Disruptions in this balance can exert dramatic consequences for neuron viability. For example, certain neuron specific RBPs HuB, HuC and HuD can enhance RNA stability by upregulating alternative polyadenylation and their loss can sensitize neurons to oxidative stress^46,47^.

### ➢ mRNA Stability Functions as a Buffering System between Transcription and Translation

There is a commonly accepted interpretation regarding the correlation of RNA stability with transcript levels: when transcription rate is stable, longer RNA half-life results in higher RNA levels^48^. Furthermore, RNAs that are constantly required at high levels are likely selected to be more stable, saving the energetic cost of transcribing and degrading transcripts. Therefore, mRNA stability (i.e., half-life) can be regarded as a predictor of transcript level or translation rate^49,50^. However, this interpretation focuses on mRNA levels within a given system, for example, a given cell, and also focuses on comparing mRNA stability between different mRNAs. In that sense, mRNAs with longer half-lives will remain longer in the RNA pool and be translated more. On the other hand, our analysis focused on comparing two systems, in which each mRNA was analyzed in either system, not compared with other mRNAs. Thus, the absolute half-lives are not at play, rather, how much did the half-life of a given mRNA change from system A to system B. While our integrative analysis of mRNA stability, transcription and translation also disclosed a positive correlation pattern in genes functionally enriched in the cell cycle related pathways. The strongest effect observed was that RNA stability countered the alterations from transcription by shifting the stability in the opposite direction. We hypothesize that this process might function to protect the translational machinery from over or under transcription of important genes, thus maintaining the functioning proteome output at physiological rates. This negative pattern was validated by measuring the gene expression, protein expression and mRNA stability of various genes. For example, WNT7A showed 15-fold increased mRNA expression in Diff. might be buffered by its destabilization, leading to only 2-fold increased protein expression. Moreover, for the key neuronal marker SYN, 3-fold increased mRNA expression in Diff. might be buffered by its destabilization, leading to 1.6-fold increased protein expression (Data not shown). Our findings are supported by a recent study in *Drosophila Melanogaster* that found that global alterations in transcriptional dynamics led cells to rapidly and specifically adjust the expression of their RNA degradation machinery in order to counteract the changes and buffer mRNA levels^23^.

### ➢ RBPs dynamically regulate mRNA stability during neuronal differentiation

SAMD4 and SRSF were the top 2 RBPs responsible for the mRNA stability changes. Previous studies have shown that Ser/Arg (SR)-rich splicing factor (SRSF) is involved in the regulation of mRNA metabolism, which has been linked to neurodevelopmental disorders^51^. However, there is no direct evidence regarding the role of SAMD4A in neuronal differentiation. SAMD4A, also known as Smaug1 in Drosophila, is a conserved RBP encoded by the SAMD4A gene^52^. Accumulating evidence highlighted the importance of SAMD4A in cell differentiation and developmental diseases^53^. For instance, depletion of SAMD4A leads to developmental defects in mice such as delayed bone development and reduced osteogenesis by inhibiting the translation of Mig6^32^. Additionally, the mouse Smaug1 protein is expressed in the central nervous system and primarily accumulates in the post-synaptic densities, which are involved in the formation of RNA granules and regulation of translation in neurons^54^. However, it was unclear to what extent findings obtained in animal model can be extrapolated to human neuronal differentiation.

This study, for the first time, shows that knockdown of SAMD4A enabled cells to retain their stemness and partially lose their ability to differentiate. Accordingly, the resistance to AntiA, Rot, Er and RSL in the SAMD4A KD cells also might be the consequence of neural stemness maintenance. This phenomenon might be attributed to the high expression of Nanog and SOX2, both of which were found to confer attenuating the adverse effects of mitochondria dysfunction and ferroptosis^55,56^. However, no difference was observed in the response to AS between SAMD4A KD and Mock cells, which re-verifies that neuronal sensitivity to stress is a selective process, and not haphazard. Furthermore, this study also provided insight into the intricate molecular mechanisms that govern the neural stemness and susceptibility to stress resulting from the knockdown of SAMD4A. SAMD4A fine-tuned the mRNA stability and expression of different transcripts individually, rather than globally enhancing/reducing stability. SAMD4A enhanced the expression of IQGAP1, a gene that promotes neuronal differentiation, but suppress the expression of PPA1, a gene that inhibits neuronal differentiation and IPO4, a gene that drives cell proliferation^57–59^. Besides, silencing SAMD4A leads to an upregulation of DENR, which acts as a protective factor against cellular stress^60^. Previous microarray-based gene expression profiling analysis showed that Smaug only destabilized targeted mRNA, as SREs strongly enriched during Smaug-dependent degradation^61^. However, this study revealed, for the first time, that SAMD4A can have a bi-directional effect on mRNA stability to regulate specific gene sets that regulate neural differentiation.

### ➢ Limitations

There are some limitations of this study that need to be acknowledged. First, the main findings are concluded based on SH-SY5Y neuronal differentiation model, which are neuronal like cells and cannot be regarded as primary neurons. Second, this study only discussed RBPs as a major regulatory factor to mRNA stability and its buffering effect as well as codon usage and optimality, while miRNAs, mRNA structures and modifications and other potential mechanisms impacting mRNA stability were not considered. Indeed, exploring all of these factors cannot be performed in a single study. However, it will be of interest to evaluate how other factors can play a role in this system as well as others. Future efforts will be imperative to analyze the various processes described here and validate their potential roles in neuronal differentiation and functioning.

## ➢ Conclusion

This study emphasizes the intricate interplay between mRNA stability, transcription, translation, and neuronal behavior. The findings suggest that mRNA stability acts as a regulatory layer to buffer against changes in transcription and translation induced by neuronal differentiation. Additionally, the identified role of SAMD4A sheds light on the intricate post-transcriptional regulatory networks governing neuronal phenotypic characteristics and stress responses. This research contributes to our understanding of the molecular mechanisms underlying neuronal differentiation and stress adaptation, providing valuable insights for future studies and potential new insights into neurodegenerative diseases.

## Materials and Methods

### ➢ Cell culture

Human SH-SY5Y neuroblastoma cells obtained from ATCC (Cat# CRL-2266) were cultured in Eagle’s Minimum Essential Medium (EMEM; ATCC, Cat# 30-2003) and Ham’s F-12 Nutrient Mixture (F12; Gibco, Cat# 11765054) 1:1 containing 10% heat-inactivated Fetal Bovine Serum (FBS; Corning, Cat# 27419002), at 37℃ and 5% CO2. No antibiotics were added to the growth media.

### ➢ Differentiation protocol

The neuronal differentiation protocol for SH-SY5Y cells was adapted from M Encinas et al ^62^. The differentiation protocol consisted of two stages: a 4-day pre-differentiation step in EMEM/F-12 supplemented with 10% FBS and 10uM RA (Sigma-Aldrich, Cat# R2625), and a subsequent 6-day differentiation step in serum-free medium containing 50 ng/ml human brain derived neurotrophic factor (BDNF) (Sigma-Aldrich, Cat# B3795). Cells were seeded at an initial density of 2 × 10^4^ cells/cm^2^ in 24-well plate coated with Type I collagen (Corning, Cat# 3524). Media were routinely changed every 2-3 days. Differentiation efficiency was evaluated by observing morphological alterations under phase contrast microscope (Leica DMi1) and immunofluorescent staining and western blot for neuronal markers and stem cell markers.

### ➢ Immunofluorescent staining

Cells seeded on 8 well glass slide (Millipore, Cat# PEZGS0816) coated with poly-L-lysine (Sigma-Aldrich, Cat# P6282) were fixed with 2% paraformaldehyde (PFA, Fujifilm Wako, Cat# 162-16065) in phosphate buffered saline (PBS) for 30 min at 4°C. Cells were permeabilizated with 0.1% Triton X-100 (Nacalai Tesque, Cat# 35501-15) for 10 min and then blocked by 2% bovine serum albumin (BSA) (Nacalai Tesque, Cat# 01860-07) for 1h at 4°C. Cells were then incubated overnight with primary antibodies Nestin (Biolegend, Cat# 839801, 1:200), MAP-2 (Santa Cruz, Cat# Sc-5359, 1:50), β-tubulin-III (Cell signaling, Cat# 2128, 1:200), or Synaptophysin (Abcam, Cat# Ab14692, 1:100) diluted in PBS containing 1% BSA and 0.1% Triton-X-100 at 4 °C. Anti-rabbit and anti-goat secondary antibodies conjugated with Alexa-Fluor 488 (Invitrogen, Cat# A11034, 1:2000) or Alexa-Fluor 555 dyes (Abcam, Cat# Ab150130, 1:2000) diluted in the same antibody buffer were incubated with the cells for 2-h at 4 °C. Cells were washed once in PBS and counterstained with 4′,6-diamidino-2-phenylindole di-hydrochloride (DAPI, Vector, Cat# H-1500). Images were taken using a laser confocal microscope (Olympus IX83) and analyzed with FLUOVIEW FV3000.

### ➢ Western blot

The undifferentiated (Undiff.) and differentiated (Diff.) SH-SY5Y cells seeded in 6 well plates were homogenized in T-PER reagent (ThermoFisher, Cat# 78510), 1% Triton X-100, 1x complete protease inhibitor (Roche, Cat# 4693116001) and 1x phosphatase inhibitor (Roche, Cat# 4906845001) on ice for 30 min. The cell lysate was then sonicated and centrifuged at 16,000g for 15min and the supernatants were extracted for protein concentration measurement by Pierce BCA protein assay (ThermoFisher, Cat# 23227). For the samples of differentiation time course, both Mock and Knockdown cells were collected at the specific time point (Day0, 1, 5, 7, 10) using the same procedure. 20-30μg protein samples were loaded into 4-20% Mini-PROTEIN TGX Precast Protein Gels (Bio-Rad, Cat# 4561096), and then were transferred to 0.2µm nitrocellulose membranes (Bio-Rad, Cat# 1704158). The membranes were blocked by 5% skim milk power (Nacalai Tesque, Cat# 31149-75) in 1x Phosphate buffered saline with Tween (PBS-T, Sigma-Aldrich, Cat# 524653), followed by overnight incubation with primary antibodies at 4 ℃. After incubation, the membranes were washed three times with PBS-T and then incubated with secondary antibody at room temperature for 1 hour. The membranes were washed three times with PBS-T again and then incubated with Pierce™ ECL Western Blotting Substrate (ThermoFisher, Cat# 32106). The protein bands were visualized by ChemiDoc MP (BioRad). Digital images were processed and analyzed using Image J. The details of the primary and secondary antibodies used in this study can be found in the Antibodies of Western Blot (**Supplementary table 2**).

### ➢ Stress response assay

To comprehensively evaluate the stress response of both undifferentiated and differentiated SH-SY5Y cells, five different stressors from three categories of stress were selected: sodium arsenite (AS) to induce nonspecific oxidative stress, Antimycin A (AntiA; respiratory complex III inhibitor) and Rotenone (Rot; respiratory complex I inhibitor) as mitochondria stressors, and Erastin (Er) and 1S, 3R-RSL3 (RSL) to induce ferroptosis stress. Each of these stressors has been reported to play a significant role in neuronal damage and the pathophysiology of neurodegenerative disorders^63^. By combining the treatment of different stress inhibitors: Ferrostatin-1 (Fer-1, ferroptosis inhibitor), Necrostatin-1 (Nec-1, necroptosis inhibitor), Ac-YVAD-cmk (YVAD, Pyroptosis inhibitor) and Ac-DVED-CHO (DEVD, apoptosis inhibitor), the main program of cell death induced by stressors could be identified.

SH-SY5Y cells were differentiated as described above on the flat bottom 96-well plates with collagen coated at 10,000 cells/well. At the endpoint of differentiation, both Diff. and Undiff. cells were pretreated with four stress inhibitors: 20μM Fer-1 (Sigma-Aldrich, Cat# SML0583), 40μM Nec-1 (Sigma-Aldrich, Cat# N9037), 10μM YVAD (Sigma-Aldrich, Cat# SML0429) and 10μM DEVD (Sigma-Aldrich, Cat# A0835) for 1 hour. Later, five different stressors: 4μM AntiA (Sigma-Aldrich, Cat# N8674), 0.2μM Rot (Sigma-Aldrich, Cat# 557368), 12μM AS (Sigma-Aldrich, Cat# S7400), 20μM Er (Sigma-Aldrich, Cat# E7781) and 2μM RSL (Sigma-Aldrich, Cat# SML2234) were added to inhibitor-pretreated-groups and stressor-only-groups for 24 hours. The control groups were treated with the same amount of growth media. Cells were then incubated with WST-8 (Nacalai tesque, Cat# 07553-44) for 4 hours at 37℃. The absorbance of formazan was measured at 450nm by Spectra Max microplate reader (Molecular Devices Spectramax 190). Each treatment had 6 replicate wells and was repeated twice (a total of 12 biological replicates). Cell viability was expressed as a percentage of the control. The stress response analysis of mock and knockdown cells also followed the paradigm mentioned above.

### ➢ RNA isolation and quality control

Cells were lysed in QIAzol Lysis Reagent (QIAGEN, Cat# 79306) for RNA extraction. RNA was extracted using the miRNeasy Kit (QIAGEN Cat# 217004) with a DNase digestion step following the manufacturer’s instructions. RNA purity and concentration were examined by Nanodrop (Thermo Fisher Scientific; Catalog# ND-ONE-W), and RNA integrity number (RIN) was assessed using RNA 6000 Nano Kit (Agilent, Cat# 5067-1511) on Agilent Bioanalyzer 2100. Samples with RNA integrity number ≥ 9 were used.

### ➢ RNA sequencing (RNA-seq)

RNA-seq libraries were prepared from differentiated and undifferentiated cells of three biological replicates using NEBNext Poly(A) mRNA magnetic isolation module (NEB, Cat# E7490) for mRNA enrichment and Ultra II directional RNA Library Prep Kit (NEB, Cat# E7760) following the manufacturer’s instruction. The quality of libraires was assessed by Agilent DNA 1000 kit (Agilent, Cat# 5067-1504)) on the Agilent Bioanalyzer 2100. The concentration of libraries was determined using NEBNext library Quant kit for Illumina (NEB, Cat# E7630). Libraries were then pooled and sequenced by Macrogen on Ilumina Hiseq X-ten platform. The sequencing was performed with 150 base-pair pair-end reads (150bp × 2) and the target depth was 50 million per sample.

### ➢ Messenger RNA stability profiling

Actinomycin D (ActD), a transcription inhibitor, is widely used in mRNA stability assays to inhibit the synthesis of new mRNA, allowing the evaluation of mRNA decay by measuring mRNA abundance^64^. Cells were treated with 5ug/ml ActD for 8 h, and then collected for RNA-seq as mentioned above.

### ➢ Ribosome profiling (Ribo-seq)

Cells were washed twice and scraped in ice-cold dPBS supplemented with 100 μg/ml cycloheximide (CHX, Sigma, Cat# C7698) and then pelleted by centrifuging at 1000 × g for 5 min. For ribosome purification, cells were lysed in 400 μL of polysome buffer (20 mM Tris-Cl pH 7.4, 150 mM NaCl, 5 mM MgCl_2_, 1 mM DTT, 100 µg/mL Cycloheximide, 1% Triton-X-100). Ribosome foot-printing was performed by adding 1.25U/μl RNase I (NEB Catalog# M0243L) to 400 μL clarified lysate and incubating samples on rotator mixer for 45min. TRIzol reagent was added, and RNA extracted using Qiagen miRNeasy kit. Ribosome protected fragments (RPFs) were selected by isolating RNA fragments of 27-35 nucleotides (nt) using TBE-Urea gel. The preparation of sequencing libraries for ribosome profiling was conducted via the NEBnext Multiplex Small RNA Library Prep Ser for Illumina according to the manufacturer’s protocol after end-repair of the RPFs using T4 PNK. Pair-end sequencing reads of size 150bp were produced for Ribo-seq on the Ilumina Hiseq X-ten system.

### ➢ Processing of the sequencing data

Quality control for Raw Fastq was performed using FastQC. Raw reads were then trimmed with Trimmomatic^65^ and adaptor sequences and low-quality read removed. Reads were aligned to the human reference genome hg38 (GRCh38.p13) using the splice aware aligner HISAT2^66^. Around 84.5%-87.5% of read pairs were uniquely mapped to the hg38 genome. Mapped reads BAM file were then counted to gene features by FeatureCounts^67^ with standard settings. Differentially expressed genes (DEGs) and differentially stabilized genes (DSGs) and differentially translated genes (DTGs) (defined as |Log_2_ FC| ≥ 1 and adj. P value < 0.05) were analyzed by Limma-Voom^68^ with normalization method TMM.

For the differential mRNA stability analysis, Diff-ActD and Undiff-ActD datasets were first normalized to their non-ActD treated counterparts individually. The changes of mRNAs in this comparison indicates changes in stability of mRNAs in relation to the total RNA pool. Next, we compared the changes in Diff. group to Undiff. group to get the differentially stabilized genes (DSG)^69^:

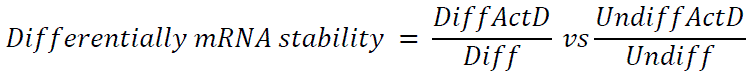

It was hypothesized that if the stability of mRNAs under the condition of undifferentiation and differentiation changed at the different scale, this will reflect on the fold change values. Genes with Log_2_FC ≥ 1 and adj. P value < 0.05 were defined as differentially stabilized genes, otherwise, genes with Log_2_FC ≤ –1 and adj. P value < 0.05 were defined as differentially destabilized genes. This analysis was performed using Galaxy^70^.

For Ribo-seq data analysis, first adapter trimming and collapsing the pair-end reads into one was done using Seqprep. Next, bowtie2 was used to align the reads to a reference of rRNA and tRNA genes to remove contaminants. After that, reads were aligned to the genome, reads counted using FeatureCounts, and differential expression conducted using Limma. Translation efficiency was calculated from RNA-seq and Ribo-seq reads using Riborex (https://github.com/smithlabcode/riborex).

### ➢ Functional Enrichment Analysis

Functional enrichment analysis and visualization were performed using pre-ranked Gene Set Enrichment Analysis (GSEA) module with the default parameters in easyGSEA in the eVITTA toolbox (https://tau.cmmt.ubc.ca/eVITTA/easyGSEA/, input date: May, 2023)^71^. Gene ontology (GO) database including Biological Process (BP), Cellular Component (CC) and Molecular Function (MF) was selected for GSEA analysis. easyVizR in the eVITTA toolbox was used to visualize the correlation among mRNA stability and transcriptome and translatome (input date: May 2023).

### ➢ Quantitative real-time PCR (qRT-PCR)

Cells were seeded in six-well plates with Collagen-I coated triplicate. After the differentiation process, both differentiated and undifferentiated cells with 5 μg/ml ActD treatment for 0h, 1h, 2h, 4h and 8h were collected at the specified time points. RNAs were extracted as described above. Total RNA concentration and purity were measured using a NanoDrop One Spectrophotometer. RNA was then converted into cDNA using the High-Capacity RNA-to-cDNA kit (Applied Biosystems, Cat# 4388950), following the manufacturer’s instruction.

QRT-PCR was performed using the GoTaq qPCR Master Mix (Promega, Cat# A6002) on a QuantStudio 5 Real-Time PCR System (Applied Biosystems), with the primers defined in the Table of Primers of qRT-PCR (**Supplementary table 3**). qPCR was conducted with 9 replicates per condition. The relative mRNA levels at each time point were analyzed using ΔΔCT method with GAPDH as the reference gene.

### ➢ Codon Analysis

A gene-specific codon counting algorithm was applied to discern the codon usage biases associated with the up– and down– stability/translated/TE genes^72^. We calculated the isoacceptor codon frequency and total codon usage frequency for genes having Log2FC > 2 or < –2 and FDR < 0.05. T-statistics values were visualized using heatmaps on Morpheus (https://software.broadinstitute.org/morpheus/). Correlation analysis at gene level was also conducted using Morpheus.

### ➢ Motif Enrichment Analysis of RNA Binding Proteins (RBPs) by Transite

The motif enrichment of RBPs was analyzed by online software; Transite (https://transite.mit.edu/)^31^. Transcript Set Motif Analysis (TSMA) and Spectrum Motif Analysis (SPMA) were performed to identify RBPs whose motifs are enriched or depleted in the DSGs dataset.

### ➢ Short Hairpin RNA design (shRNA) and knockdown experiment

Short hairpin RNA (shRNA)-encoding pairs of oligonucleotides with targeting sequences to SAMD4A and SEPHS2 mRNA was designed as follows:

SAMD4A: 5’TGCAACAGGAATCCAAGGATAATTCAAGAGATTATCCTTGGATTCCTGTTGCTT TTTTC-3’

SEPHS2: 5’TCATTGACAAGCCGCGAGTTATTTCAAGAGAATAACTCGCGGCTTGTCAATGTT TTTTC

Mock: 5’TGAAATACTCAGCAGATCATTATTCAAGAGATAATGATCTGCTGAGTATTTCTT TTTTC-3’

Each sequence was cloned into the corresponding sites of pLB vector (Addgene, Cat#11619). Lentiviruses were generated by co-transfection of Lenti-X 293T (Takara, Cat# 632180) cells with three plasmids: a lentiviral vector plasmid, pMD2.G (expressing envelop protein, FASMAC) and psPAX2 (expressing packaging proteins, FASMAC). Media were changed 16 h after transfection, and the supernatants were harvested 48 h after transfection. Cell debris in the media was removed by 0.45 µm filtration following centrifugation at 1500 g for 10 min. Viral particles were collected twice 48 h and 72 h post-transfection. For infection, lentivirus particles were added to each well of a six-well plate containing 7.5 × 10^5^ cells. Cells were incubated with lentivirus and 4 µg/ml polybrene (Sigma, Cat#TR-1003) for 12 h. The expression of Green Fluorescent Protein (GFP) was checked under the immunofluorescence microscopy. The transfection efficiency of cells was set to 100% based on fluorescence distribution and western blot was used to evaluate the knockdown efficiency.

### ➢ Neurite Growth Rate Analysis

The analysis of neurite growth rate was conducted using the images obtained from the microscope (Leica DMi1). Image J was utilized to track the length of neurites (Ridge Detection) and the surface area of cells, including both neurites and cell bodies. The ratio of neurite length to cell surface area was calculated as the percentage of neurite outgrowth activity.

### ➢ Statistical Analysis

Statistical tests were performed with GraphPad Prism 7 software. The values were presented as mean ± standard deviation (SD) of at least 3 biological replicates or as indicated. For each dataset, the Shapiro-Wilk normality test was applied to determine if the data had a normal distribution. If the data passed the normality test, the parametric test was used, otherwise the non-parametric test was used. For the analysis of western blot data, comparisons between Diff. and Undiff. / SAMD4A KD and Mock were performed with unpaired Student t-test (two-tailed). For the analysis of stress response assay and mRNA stability by qRT-PCR and neurite growth rate, two-way analysis of variance (ANOVA) with Bonferroni post-hoc test was performed. The statistical significance was set at P < 0.05 (*P < 0.05, **P < 0.01, ***P < 0.001 and ****P < 0.0001).

## Data availability

The raw sequencing data files are available through Sequence Read Archive (SRA) database with accession number PRJNA779467, PRJNA1004177, and PRJNA1001994.

## Author contribution

**YZ**: Study design. Performed all experiments. Data analysis and interpretation. Wrote the manuscript. Funding Acquisition. **SR**: Conception. Study Design. Data analysis and interpretation. Revised the manuscript. Administration. Study supervision. Funding acquisition. **TT**: Critically revised the manuscript. **KN**: Critically revised the manuscript. Study supervision.

## Acknowledgment

The authors report no conflict of interest nor there are any ethical adherences regarding this work. The authors would like to that Dr Thomas J Begley for providing the codon analysis algorithm used in this study.

