## Supplemental data for "Dynamic mRNA stability changes buffer transcriptional activation during neuronal differentiation and are regulated by RNA binding proteins"

| | Most Enriched Hexamers | | | Most Depleted Hexamers | | |
|---|---|---|---|---|---|---|
| | Hexamer | Enrichment value | adj. P | Hexamer | Enrichment value | adj. P |
| 1 | AGCGCG | 2.96877 | 2.11E-15 | GCGAUU | 0.438967 | 0.007861 |
| 2 | CGCCGG | 2.695272 | 1.94E-19 | GUAGUC | 0.45958 | 1.41E-05 |
| 3 | CCGCGA | 2.679451 | 3.08E-09 | AAGCGA | 0.540728 | 0.009492 |
| 4 | CCGCCG | 2.66048 | 1.86E-32 | UAGCUA | 0.54353 | 8.37E-05 |
| 5 | CGUCGG | 2.581627 | 1.74E-06 | GCUAAU | 0.560165 | 3.94E-06 |
| 6 | GCGCCG | 2.569668 | 1.94E-19 | GUAGCU | 0.590854 | 8.91E-06 |
| 7 | CGCGAC | 2.559722 | 2.06E-05 | GUAAUC | 0.604679 | 3.77E-05 |
| 8 | UGCGCG | 2.523651 | 5.46E-09 | CCUUAC | 0.609744 | 0.00027 |
| 9 | CGCCGC | 2.480434 | 7.55E-27 | UAGUCC | 0.612407 | 0.002477 |
| 10 | CCGACG | 2.476833 | 1.27E-07 | GCGAUU | 0.438967 | 0.007861 |

**Supplementary Table 1:** Top 10 hexamers statistically significantly (adj. P < 0.01) enriched and depleted in stabilized genes after the SH-SY5Y differentiation, related to Figure 5.

| Antibody | Manufacturer | Catalog No. | Dilution |
|---|---|---|---|
| *Primary antibody* | | | |
| Nestin | Invitrogen | MA1-110 | 1:500 |
| MAP2 | Sigma | M1406 | 1:500 |
| NF-M | Millipore | AB1987 | 1:500 |
| TUJ | Cell Signaling | 2128 | 1:1000 |
| SYN | Invitrogen | MA5-14532 | 1:1000 |
| SOX2 | Cell Signaling | 2748 | 1:500 |
| Nanog | Abcam | AB173368 | 1:1000 |
| beta-actin | Cell Signaling | 4970 | 1:1000 |
| *Secondary antibody* | | | |
| Rabbit IgG HRP | Cell Signaling | 7074 | 1:1000 |
| Mouse IgG HRP | Cell Signaling | 7076 | 1:1000 |

**Supplementary Table 2**: The Antibodies of Western Blot. Notes: Catalog No. (Catalog Number); MAP2 (Microtubule Associated Protein 2); NF-M (Neurofilament Middle); TUJ (beta-Tubulin-III); SYN (Synaptophysin); IgG (Immunoglobulin G); HRP (Horseradish peroxidase).

| Gene | Accession Number | Forward Primer | Reverse Primer |
|---|---|---|---|
| SNAI1 | NM_005985.4 | CCAGTGCCTCGACCACTATG | TTAGAGTCCTGCAGCTCGCTGTA |
| ID2 | NM_002166.5 | CAACACGGATATCAGCATCCTGTC | GCCACACAGTGCTTTGCTGTC |
| ADM | NM_001124.3 | GGTGACACTGGATAGAACAGCTCAA | GTACATCAGGGCGACGGAAAC |
| ID1 | NM_181353.3 | GCTCTACGACATGAACGGCTGTTA | GCTCCAACTGAAGGTCCCTGA |
| WNT7A | NM_004625.4 | GAGATCAGGTCTCAGGCATGG | CTTGGCGGTTCAACACAATCAC |
| SHISA2 | NM_001007538.2 | GGATTGCCTTTCTTCTCCCACA | GCAGTTGGAAATGTACTTGCACATC |
| GPR83 | NM_001330345.2 | CCATGACTTTCATTGGGCAGAC | GGCAGCTTTCAGATGAAACCATTAG |
| ARHGAP20 | NM_001258415.2 | TTGCTTCCAGACGTTGTTCCATA | ACGAGATGTACTGCTGCAACCTTTA |
| SEPHS2 | NM_012248.4 | GGATCGTTGGCATTGTGGAA | GAGCAAGAACAGCAGCTGTGG |
| IQGAP1 | NM_003870.4 | TGGGCAATTCTGTTTGTGTAACTC | AGGTGCCTTGACCCATTCTG |
| PPA1 | NM_021129.4 | ATGTTGCGAATTTGTTCCCGTA | TCATTGTCACCACAACAGCCAGTA |
| DENR | NM_003677.5 | GGTGAAGCAGATTTATCCATCGAGA | TCACATGTTGATACAGGCACACAGA |
| IPO4 | NR_051979.2 | AGCTCTGTGGCGTGCTCAAG | GGCGTCGTATTCAGCCTGTG |
| GAPDH | NM_002046.7 | GCACCGTCAAGGCTGAGAAC | TGGTGAAGACGCCAGTGGA |

**Supplementary Table 3:** Primers of qRT-PCR.

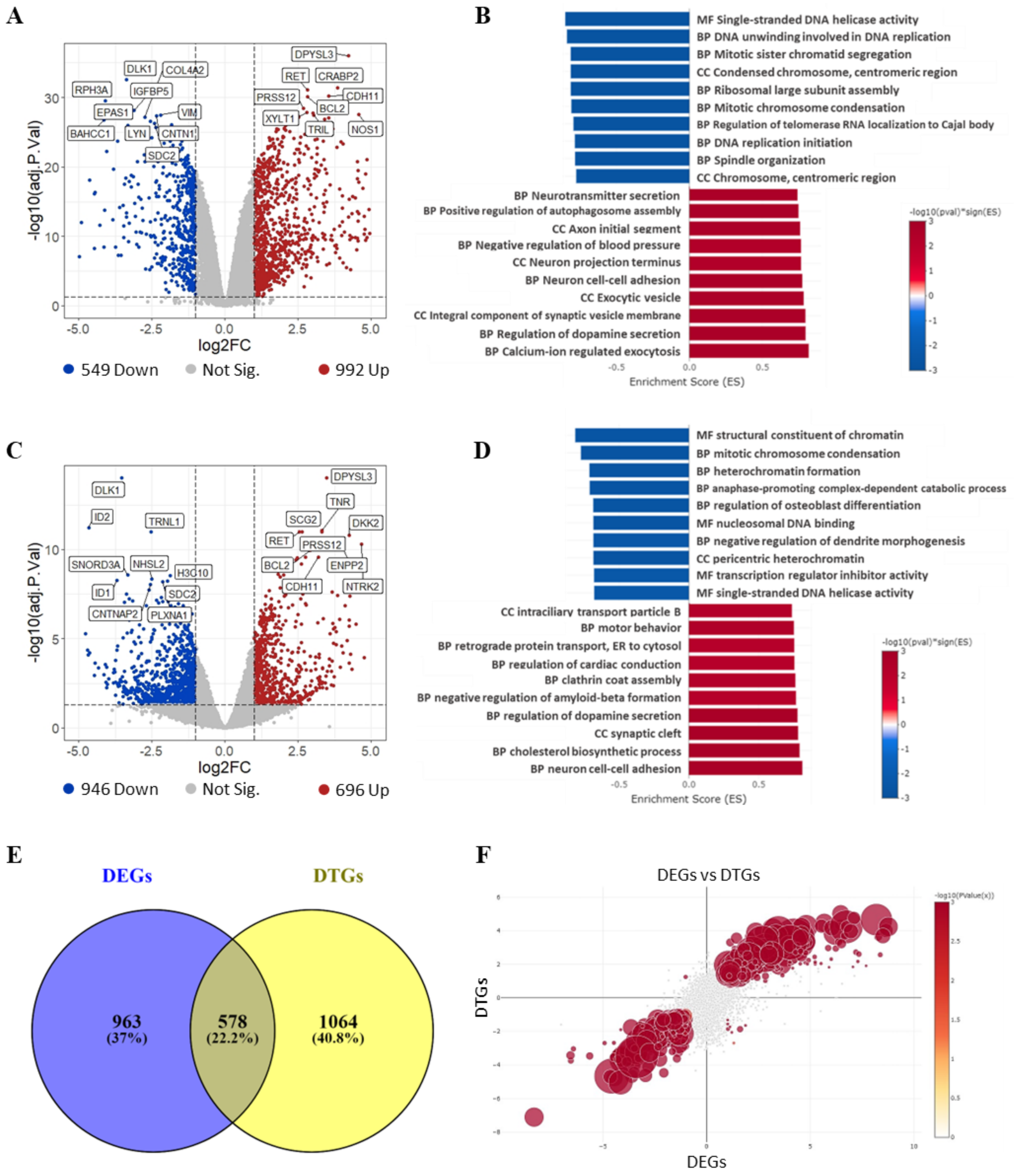


**Supplementary Figure 1** Transcriptomics and translatomics and their interplay in the differentiated SH-SY5Y cells. **A** The volcano plot showing the $\log_2$ FC and $-\log_{10}$ adj. P value of each gene. Significantly up and down differentially expressed genes (DEGs) were indicated by red and blue dots, respectively. (Cutoff: $|Log_2FC| \geq 1$ and adj. $P < 0.05$). Top 10 up and down DSGs with the lowest adj. P value were labeled by gene symbol. **B** GSEA identifying Top 10 enriched GO pathways of up- (Red) and down-regulated genes (Blue), respectively. **C** The volcano plot comparing the translational $\log_2$FC and $-\log_{10}$ adj. P of genes in differentiated vs undifferentiated cells. Significantly upregulated and downregulated genes were marked in red and blue, respectively. D Bar graph showing Top 10 enriched gene ontology terms of up- (Red) and down-regulated genes (Blue), respectively. **E** The Venn diagram depicting the overlap between DEGs and DTGs. **F** The scatter plot showing the Pearson correlation between DEGs and DSGs. Correlation $R^2 = 0.909$.

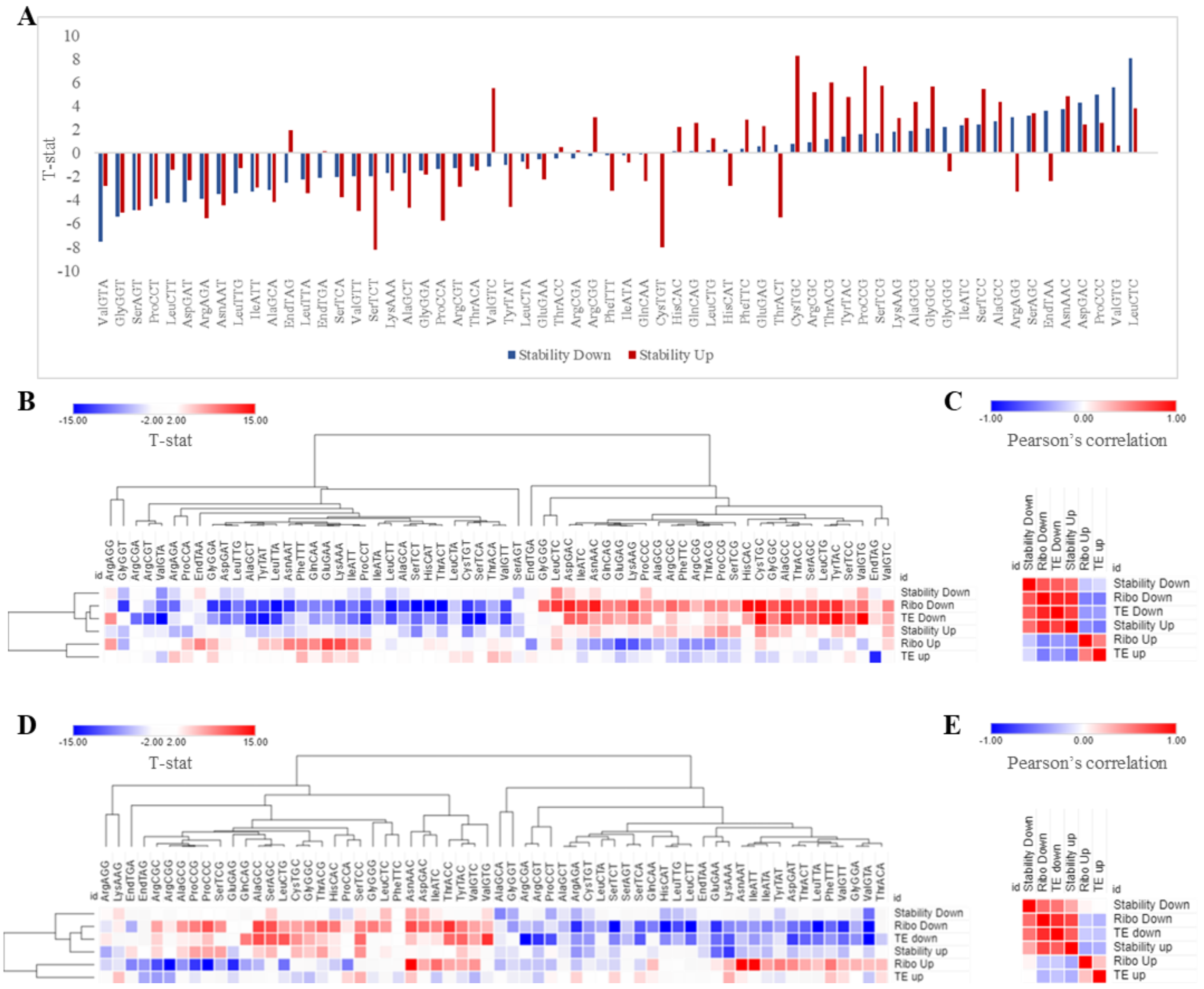


**Supplementary Figure 2**: Codon usage of RNA stability, Translatome and Translation Efficiency (TE, Translatome vs Transcriptome) in the differentiated SH-SY5Y. **A** The codon occurrence to RNA stability potted for each codon. **B** Heatmap of synonymous codon usage in the dataset of stability, translatome and TE. Up indicating differentially stabilized genes, up-translated genes and higher TE genes, while down indicating differentially destabilized genes, down-translated genes and lower TE genes. **C** Pearson’s correlation coefficient analysis among each dataset. **D** Heatmap of global codon usage in the dataset of stability, translatome and TE. Up indicating differentially stabilized genes, up-translated genes and higher TE genes, while down indicating differentially destabilized genes, down-translated genes and lower TE genes. **E** Pearson’s correlation coefficient analysis among each dataset.

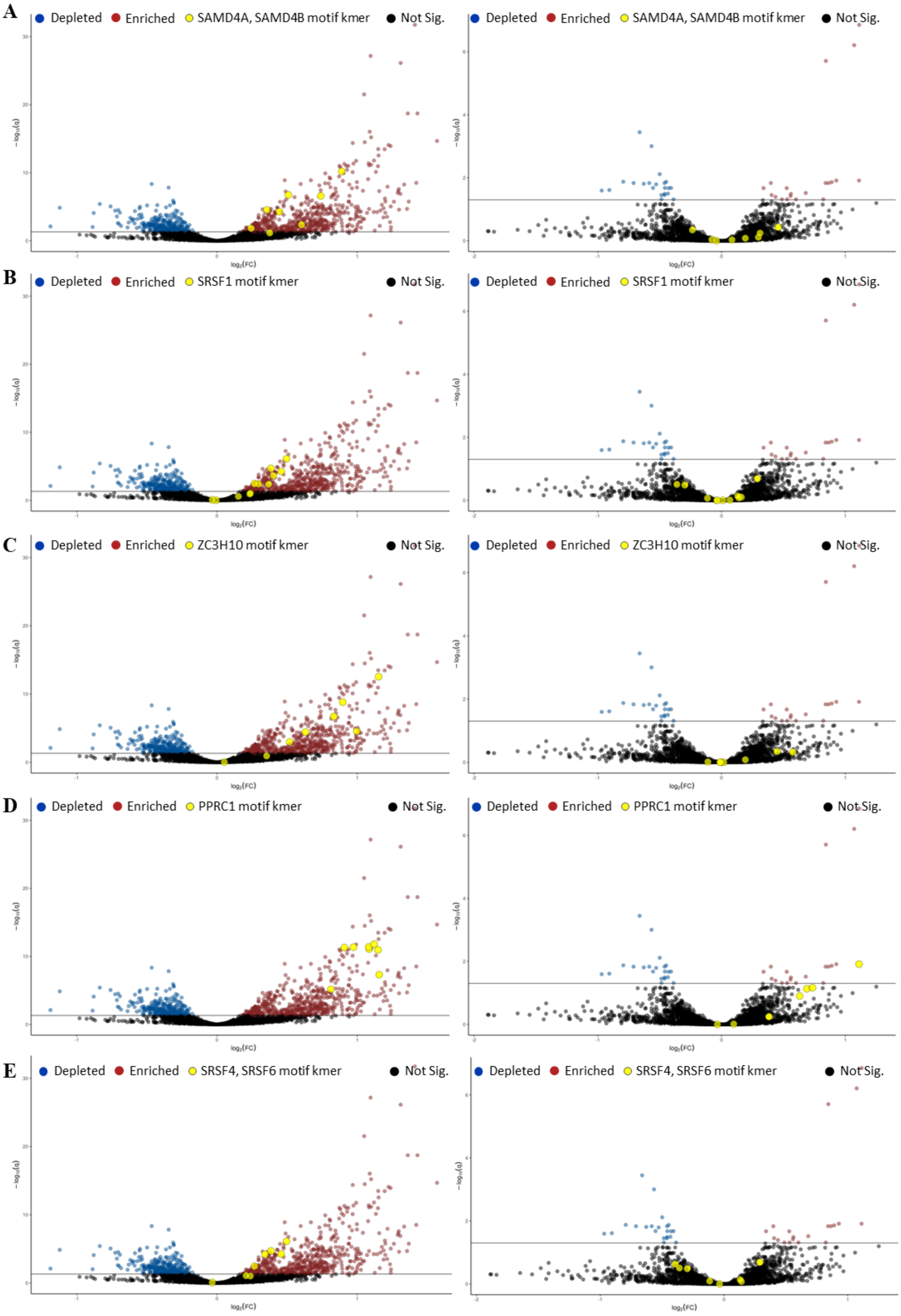
A
Depleted
Enriched
SAMD4A, SAMD4B motif kmer
Not Sig.
B
Depleted
Enriched
SRSF1 motif kmer
Not Sig.
C
Depleted
Enriched
ZC3H10 motif kmer
Not Sig.
D
Depleted
Enriched
PPRC1 motif kmer
Not Sig.
E
Depleted
Enriched
SRSF4, SRSF6 motif kmer
Not Sig.
$-\log_{10}(q)$
$\log_2(FC)$

**Supplementary Figure 3** Volcano plot of Top5 RBPs identified by k-mer-based TSMA, related to **Figure 5**. **A** TSMA volcano plot showing enriched and depleted k-mers of SAMD4A, SAMD4B in stabilized transcripts (left panel) and destabilized (right panel). **B** TSMA volcano plot showing enriched and depleted k-mers of SRSF1 in stabilized transcripts (left panel) and destabilized (right panel). **C** TSMA volcano plot showing enriched and depleted k-mers of ZC3H10 in stabilized transcripts (left panel) and destabilized (right panel). **D** TSMA volcano plot showing enriched and depleted k-mers of PPRC1 in stabilized transcripts (left panel) and destabilized (right panel). **E** TSMA volcano plot showing enriched and depleted k-mers of SRSF4, SRSF6 in stabilized transcripts (left panel) and destabilized (right panel).

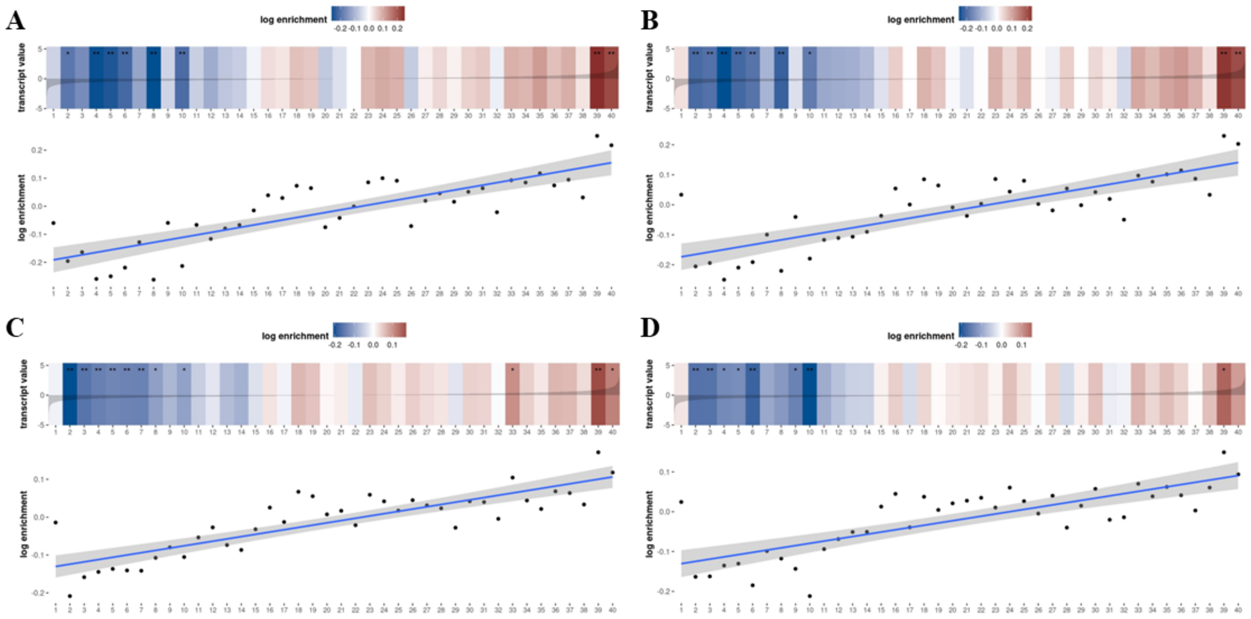


**Supplementary Figure 4** Top spectrum plots of matrix based SPMA, related to Figure 5. **A** Enriched in stabilized genes: M154 0.6 – SRSF1. **B** Enriched in stabilized genes: M126 0.6 – SRSF1. **C** Enriched in stabilized genes: M065 0.6 – SRSF9. **D** Enriched in stabilized genes: 359 12507992.

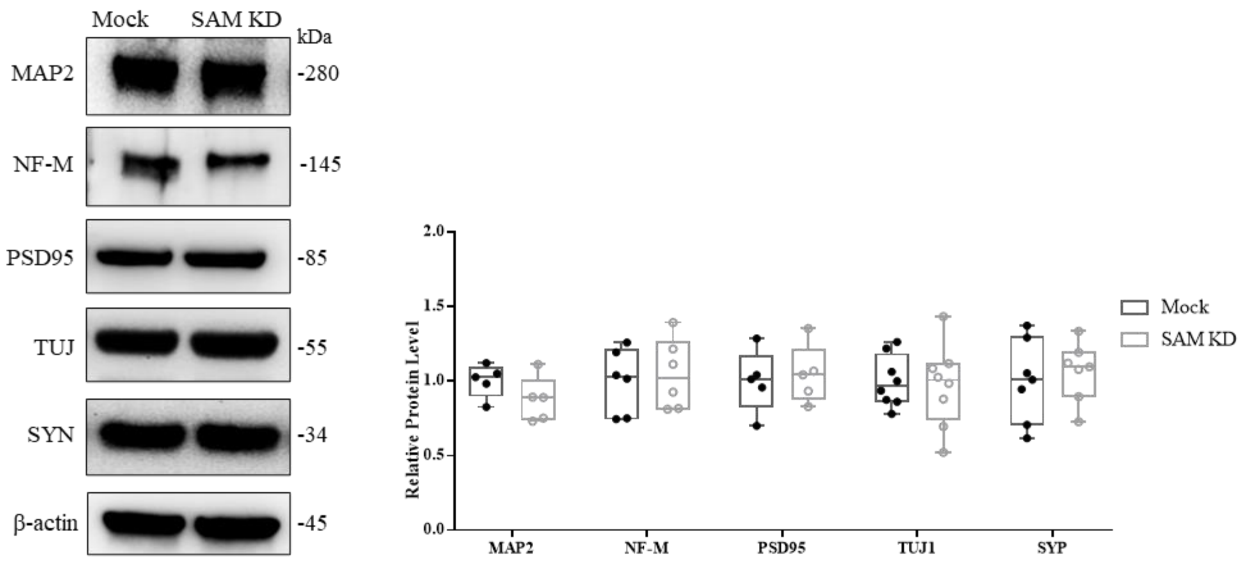


**Supplementary Figure 5** The expression of neuronal markers in SAMD4A Knockdown and Mock Cells, band intensity quantified by ImageJ.

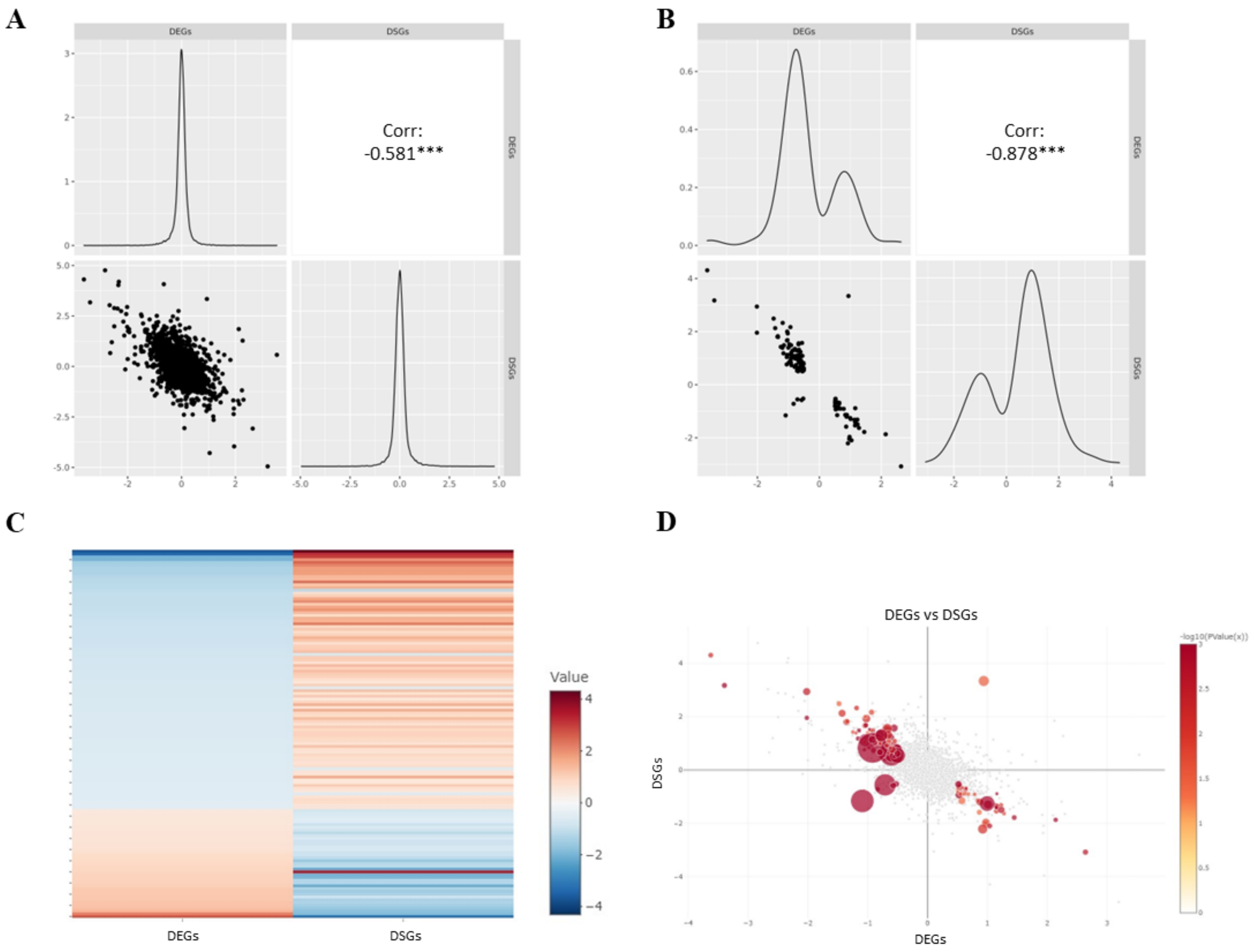


**Supplementary Figure 6** mRNA Stability Negatively Correlates with Transcriptomics in SAMD4A KD cells. **A** Correlation analysis of mRNA stability and transcriptome involving all the transcripts in SAMD4A. **B** Correlation analysis among DEGs and DSGs with applying cutoff: $|Log_2FC| \geq 0.5$ and $P < 0.05$). **C** Heatmap showing $Log_2FC$ of DEGs and DSGs. **D** Visualization of the $Log_2FC$ values of each overlapped genes between DSGs and DEG.
